# METTL3 Uncouples Chromatin Accessibility from Transcription during Retinal Development

**DOI:** 10.1101/2025.10.02.680012

**Authors:** Jing Xu, Yuanhao Huang, Zhaowei Han, Qiang Li, Jie Liu, Rajesh C. Rao

**Author notes:** Corresponding author: Rajesh C. Rao, MD University of Michigan Medical School W.K. Kellogg Eye Center 1000 Wall St. Brehm Rm 8333 Ann Arbor, MI 48105. Authors contributed equally (Jing Xu, Yuanhao Huang).

## Abstract

METTL3 is a key regulator of RNA metabolism, yet its genomic and epitranscriptomic roles in tissue development are largely unexplored. Using embryonic stem cell-derived 3D retinal organoids to model retinal progenitor cell (RPC) differentiation, we integrated transcriptome-wide m6A profiling (GLORI), protein-DNA (ChIP-seq and CUT&RUN) and chromatin accessibility (ATAC-seq) mapping, and targeted m6A engineering (dCas13b-FTO) to dissect METTL3 function. Loss of METTL3 nuclear m6A activity disrupted *Rx*+ retinal anlage formation *in vitro*, with dCas13b-FTO epitranscriptome engineering revealing that m6A at the *Six3* 3’UTR governs its stability. Surprisingly, while METTL3 loss altered histone modifications and chromatin accessibility, its direct chromatin targets showed little transcriptional correlation. A degron-based METTL3 degradation strategy, paired with protein-RNA interaction profiling, exposed rapid regulatory shifts in RPCs, revealing a METTL3-*Ythdf1* protein-RNA axis. Our multi-omics approach establishes METTL3-dependent m6A as a critical epitranscriptomic layer in retinal development, unveiling a genomic paradigm in which chromatin accessibility diverges from transcriptional output.

## Introduction

20-40% of all mRNAs are methylated (Kim, van Galen et al. 2021). Yet, our understanding of RNA methylation (epitranscriptomics) and its crosstalk with the epigenome in tissue development is limited. Discovery of epigenetic factors modifying chromatin have revealed pathways upstream of eye field transcription factors (EFTFs), which are required for retinal development. For instance, we found that WDR5, a histone methyltransferase co-factor, controls EFTF activity and cell fate during retinogenesis (Li, Mao et al. 2020, Li, Huang et al. 2021). RNA methyltransferases (“writers”) such as METTL3 and METTL14 associate in an *N*^6^-methyladenosine (m6A) methyltransferase complex to modify RNA, which regulates RNA metabolism, including stability (Liu, Yue et al. 2014).

Despite the relative abundance of m6A RNA methylation, our understanding of the role of *Mettl3* and the function of m6A modifications in retinal development is incomplete. In zebrafish, *Mettl3* knockdown leads to microphthalmia due to delayed retinal progenitor cell (RPC) differentiation and increased cell death (Huang L 2021). In mice, *Mettl3* loss leads to abnormalities in RPC-to-Müller glia transition in late retinogenesis due to increased half-life and reduced m6A of RPC transcripts (Xin, He et al. 2022). Similar findings were reported for *Mettl14* inactivation and triple deletion of m6A readers *Ythdf1*, *Ythdf2*, and *Ythdf3* (Niu, Che et al. 2022, Li, Sun et al. 2023). *Zfp292*, which inhibits cell cycle progression, showed the highest reduction in m6A in both SIX3^+^ RPCs with *Mettl3* deletion and CHX10^+^ RPCs with *Mettl14* loss. Overexpression of *Zfp292* phenocopied the effects of *Mettl3* and *Mettl14* inactivation in late RPCs (Xin, He et al. 2022, Li, Sun et al. 2023). These reports are the first foray into retinal epitranscriptomics, but questions remain. To date, no study has explored the direct RNA targets of METTL3, nor its chromatin targets and whether epitranscriptomics-epigenetics crosstalk occurs during retinal development.

Prior work has focused on canonical effects of METTL3 on RNA modifications, but few studies have interrogated the broader epigenomic impact of METTL3 in tissue development. Here, we use the pluripotent cell-to-*Rx+* eye field transition—a pivotal stage of retinal anlage formation—by leveraging mouse embryonic stem cell (mESC)-derived retinal organoids and *Mettl3*-deficient *in vivo* models to probe METTL3 function across two phases: (1) mESC-to-RPC and (2) RPC-to-retina differentiation. Using state-of-the-art epigenomic tools—genome-wide m6A mapping (GLORI), METTL3-RNA profiling, dCas13b-FTO m6A engineering, PROTAC-based degradation, and integrative NGS (CUT&RUN)— we reveal the primacy of catalytic activity of nuclear METTL3, the functional role of 3’UTR m6A modifications in regulation of an EFTF RNA, *Six3*, a regulatory METTL3-*Ythdf1* protein-RNA interaction –modes of epitranscriptomic control that influence acquisition of the retinal cell fate. We observe that METTL3 binds RPC chromatin, regulates bivalent histone modifications and chromatin accessibility, but these events do not alter transcription. Our work uncovers a METTL3-mediated uncoupling of chromatin accessibility and transcription that occurs in the context of METTL3 control of RNA stability during development of critical tissue: the retina.

## Results

### Enzymatic function of nuclear METTL3 controls the mouse embryonic stem cell-to-retinal progenitor cell (mESC-to-RPC) fate transition

The five-to-six-day duration mESC-to-mouse retinal organoid (mRO) differentiation platform is a rapid, powerful system to interrogate mechanisms that regulate the transition from pluripotency to formation of the eye field and optic vesicle, the earliest events of retinal development. The related molecular events cannot be easily studied *in vivo* due to the dearth of isolable tissue that limit NGS-based analyses. We leveraged this system to first survey the m6A landscape and its potential importance in differentiation of Rx+ RPCs, the predominant cell type in mROs. To this end, we use an mESC line containing GFP knocked into one allele of *Rx* gene, which encodes a key eye field transcription factor EFTF required for RPC specification. By m6A nanopore-based sequencing (Liu, Begik et al. 2019), we mapped, genome-wide, the m6A landscape in day 6 RPCs. The distribution of m6A sites (n= 26,729) within transcripts was mainly at 3’UTR (56.33%) and CDS (∼40%), with only about 4% of m6A sites detected localize at 5’UTR (**Fig S1A**), consistent with previous reports in other cell types (Dominissini D 2012, Meyer KD 2012, Linder B 2015). METTL3 is the primary RNA methyltransferase that catalyzes m6A deposition. To interrogate its role, we generated Rx:GFP *Mettl3* rescue complementation mESC lines, wherein a doxycycline (Dox)-inducible form of ectopic *HA*-tagged *Mettl3* was introduced via a Piggybac transposon system (HA-METTL3^WT^), and endogenous *Mettl3* was inactivated by CRISPR-Cas9 editing (*Mettl3^KO^*), through a strategy we previously reported (**Fig 1A)** (Li Q 2020). By analysis of independent *Mettl3^KO^*;HA-METTL3^WT^ Rx:GFP mESC clones, we found that *Mettl3^KO^* mESCs were unable to form Rx+RPCs. Rescue of Rx:GFP+ RPC differentiation was seen with expression of Dox-inducible HA-METTL3^WT^, indicating that *Mettl3* is required for proper Rx+ RPC differentiation (**Fig S1B** and **Fig 1B**).

**Figure 1.**
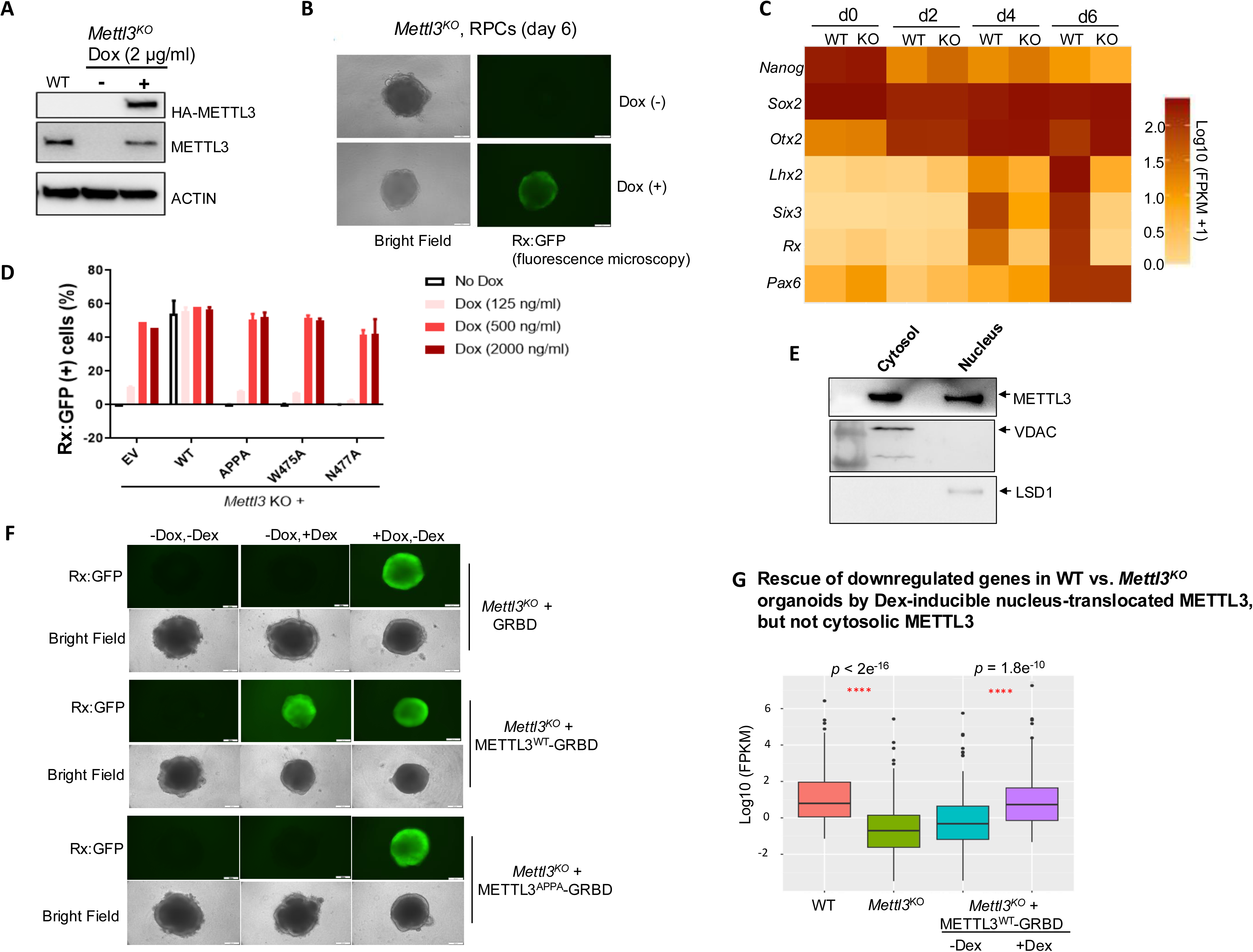
Nuclear METTL3 with intact catalytic activity is required for the mESC-to-RPC cell fate transition. (**A**) Generation of *Mettl3^KO^* Rx:GFP mESCs with ectopic Dox-inducible HA-METTL3^WT^ rescue. Endogenous *Mettl3* deletion and exogenous METTL3^WT^ re-constitution were detected via western blot using METTL3 and HA antibodies, respectively. ACTIN served as a loading control, n=3 independent experiments. (**B**) *Mettl3*^KO^ mESCs displayed impaired Rx:GFP+ RPC differentiation at day 6 (d6), n=3 independent experiments. (**C**) Time course heatmap of mESC pluripotency and eye field transcription factor transcripts under RPC culture conditions during WT and *Mettl3*^KO^ mESC differentiation, n=3 independent experiments. (**D**) METTL3 catalytic activity is required for proper Rx:GFP+ RPC differentiation. WT and catalytic mutant METTL3 plasmids (APPA, W475A, N477A) were introduced into *Mettl3*^KO^ mESCs and cultured in Rx:GFP+ RPC differentiation conditions. Day 6 Rx:GFP induction by Dox dose dependent HA-METTL3^WT^ in the presence of empty vector (EV), WT and mutant FLAG-METTL3 was quantified by flow cytometry, n=3 independent experiments. (**E**) METTL3 protein is present in both cytosol and nucleus of Rx:GFP+ RPCs at day 6. Rx:GFP+ RPCs were sorted by flow cytometry, followed by cell fractionation and western blotting with METTL3, VDAC and LSD1 antibodies. VDAC and LSD1 served as cytosolic and nuclear markers, respectively, n=3 independent experiments. (**F**) Cytosol-to-nuclear METTL3 translocation was induced by adding Dex (10 μM), which rescued the Rx:GFP+ expression in *Mettl3*^KO^ RPCs, but cytosolic METTL3 (without Dex, without Dox) did not. As a positive control, Dox-inducible METTL3^WT^ (+Dox at 2 μg/ml) in *Mettl3*^KO^ mESCs also rescued Rx:GFP+ RPC differentiation. After 6 days under Rx:GFP RPC differentiation conditions, organoids were visualized by bright field and fluorescence microscopy. *Mettl3*^KO^ reconstituted with GRBD alone also served as negative control. Scale bar: 200 μm, n=3 independent experiments. (**G**) Bulk RNA-seq showed differentially expressed genes in *Mettl3*^KO^ organoids were rescued by nucleus-translocated METTL3 (+Dex), but not cytosolic METTL3 (-Dex). Downregulated genes in WT vs *Mettl3*^KO^ RPCs and Dex-rescue (+Dex vs -Dex) (day 4) were compared by pairwise student t-tests (asterisks denote p < 0.05). All other pairwise comparisons were non-significant (p > 0.05), n=3 independent experiments

To determine whether retinal differentiation defects due to *Mettl3* loss are lineage-specific and/or a result of impaired exit from pluripotency, in our organoid culture platform, we performed RNA-seq every 2 days over the mESC-to-RPC 6-day differentiation period (**Fig 1C)** and validated this with RT-qPCR (**Fig S1C**). In both WT and *Mettl3^KO^* organoids. We observed comparable loss of pluripotency gene *Nanog* as well as similar induction of *Sox2*+/*Otx2*+/*Pax6+* neuroectodermal genes in both WT and *Mettl3^KO^* organoids. In contrast, the induction of EFTF genes *Lhx2*, *Six3* and *Rax* was remarkably impaired (**Fig 1C** and **Fig S1C**), consistent with no detectable Rx:GFP+ RPC differentiation in *Mettl3*^KO^ ESCs (**Fig S1B** and **Fig 1B).** Thus, *Mettl3* appears to have a stronger effect on retinal neuroectoderm differentiation vs. that of non-retinal neuroectoderm from mESCs.

We next asked whether the defect in retinal organoid differentiation in *Mettl3^KO^* mESCs is dependent on METTL3 catalytic activity. To this end, we mutated the conserved METTL3 catalytic motif from DPPW to APPA to abolish METTL3 enzymatic activity (Wang P 2016). Also, W475A and N477A mutants dramatically reduce its catalytic activity via disruption of the interaction between METTL3 and METTL14 (Wang P 2016). We constitutively expressed the FLAG-tagged METTL3^WT^ and METTL3^APPA^, METTL3^W475A^, METTL3^N477A^ mutants into *Mettl3^KO^* Rx:GFP mESCs (**Fig S1D**). None of the catalytic mutants rescue the defective RPC differentiation in *Mettl3^KO^* mESCs, in contrast FLAG-METTL3^WT^ which rescued RPC formation (**Fig 1D**). Thus, METTL3 catalytic activity is required for proper RPC formation.

Since METTL3 is present in both the cytosol and nucleus during mESC-to-RPC differentiation (**Fig 1E**), we asked whether compartment-specific METTL3 is required for RPC differentiation. To this end, we constitutively expressed FLAG-tagged, GRBD, METTL3^WT-GRBD^ and METTL3^APPA-GRBD^ into *Mettl3*^KO^;*HA-METTL3^WT^* Rx:GFP mESCs. In this system METTL3-GRBD fusion proteins are translocated from the cytosol to nucleus by dexamethasone (Dex) culture addition (Murakami, Gunesdogan et al. 2016).

Western blot with FLAG antibody (Sigma) showed the equivalent level of FLAG-tagged METTL3^WT-GRBD^ and METTL3^APPA-GRBD^ with or without Dex (10μM) treatment (**Fig S1E**). Nucleic METTL3^WT-GRBD^ (+Dex), but not the cytosolic METTL3^WT^ (-Dex; -Dox), rescued the Rx:GFP+ RPC differentiation from *Mettl3^KO^* mESCs (**Fig 1F, Fig S1F**). Neither nucleic METTL3^APPA^ (+Dex) nor cytosolic METTL3^APPA^ (-Dex; -Dox) could rescue the defects (**Fig 1F, Fig S1F**). mRNA-seq showed that the dysregulated differentially expressed genes (DEGs) in *Mettl3^KO^* RPCs were rescued by nucleic METTL3 but not cytosolic METTL3 (**Fig 1G, Fig S1G**). Collectively, reconstitution with METTL3 catalytic mutants and forced cytosolic localization of METTL3 were not able to induce RPC differentiation from *Mettl3*^KO^ mESCs, indicating that defective mESC-to-RPC differentiation requires nuclear METTL3 catalytic activity.

### METTL3-binding to chromatin is uncoupled from transcription in mESC-derived RPCs

METTL3 non-canonically regulates chromatin accessibility in mESCs via an m6A-dependent interaction with histone modifying enzymes and chromatin regulatory RNAs, as well as the senescence associated secretary phenotype in fibroblasts through m6A-independent association with enhancers (Liu, Dou et al. 2020, Liu, Li et al. 2021, Xu, Li et al. 2021). Moreover, bivalent chromatin dynamics accompanies early mESC lineage commitment (Ruthenburg AJ 2007, Schwartz YB 2007, Liu X 2016). We wondered whether similar non-canonical METTL3 functions may be involved in the mESC-to-RPC fate transition. To this end, we mapped genome-wide H3K4me3 and H3K27me3 distribution, which are associated with gene activation and repression respectively, and performed parallel RNA-seq, in WT and *Mettl3*^KO^ RPCs at day 4 by CUT&RUN (Li, Mao et al. 2020, Li, Huang et al. 2021). We observed *Mettl3*-dependent H3K4me3 marks that were not only depleted of this modification in *Mettl3*^KO^ RPCs, but enriched with the repressive H3K27me3 modification at these sites (**Fig 2A**). When we integrated RNA-seq with CUT&RUN, we found that the down-regulated genes in *Mettl3* KO vs WT RPCs were indeed less enriched with H3K4me3 modifications, but these sites harbored H3K27me3 marks (**Fig 2B**). Our data indicates METTL3 associates with specific H3K4me3 modifications in *Mettl3*-dependent DEGs, likely related to activation of specific genes during the mESC-to-RPC differentiation program.

**Figure 2.**
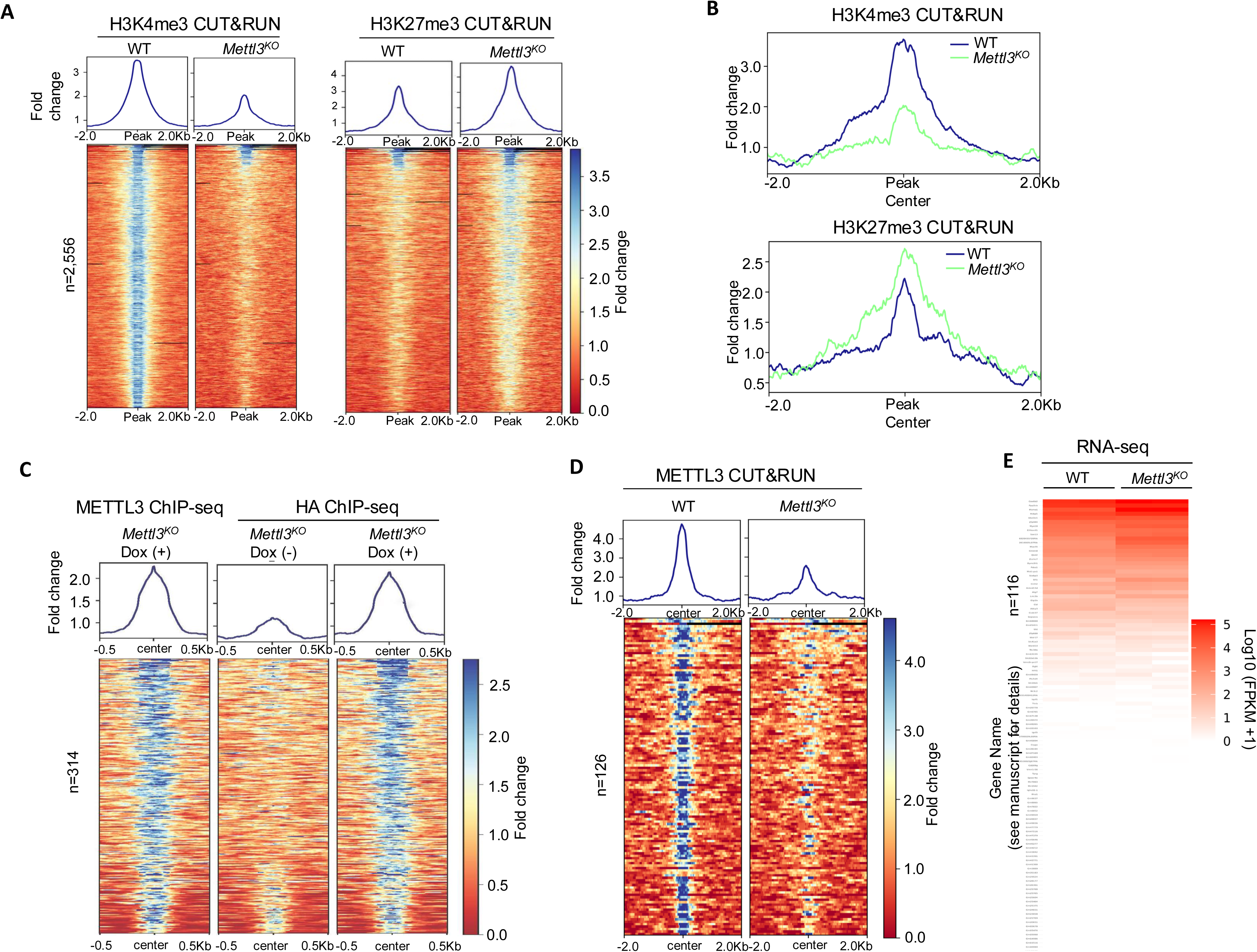
METTL3 regulates histone modifications but its direct targets on chromatin do not contribute to gene regulation during mESC-to-RPC differentiation. (**A**) Heatmap showing fold change of H3K4me3 and H3K27me3 enrichment in WT and *Mettl3*^KO^ organoids under RPC differentiation conditions at day 4, n=3 independent experiments. (**B**) Downregulated genes in *Mettl3^KO^* organoids under RPC differentiation conditions (i.e., day 4 group in Fig.1C) showed reduced H3K4me3 and increased H3K27me3 recruitment. (**C**) Heatmap showing fold change of HA and METTL3 antibodies ChIP enrichment in *Mettl3*^KO^ organoids reconstituted with exogenous Dox-inducible METTL3^WT^-HA after 6 days in RPC differentiation conditions. (**D**) Heatmap showing fold change of METTL3 antibody CUT&RUN enrichment in WT and *Mettl3^KO^* organoids after 4 days in RPC differentiation conditions, n=3 independent experiments. (**E**) Heatmap for gene expression at METTL3-bound genes (integration of **Figs. 1C and 2D** RNA-seq datasets in WT and *Mettl3^KO^* organoids after 4 days in RPC differentiation conditions), n=3 independent experiments.

To identify potential METTL3 targets on chromatin, we used ChIP-seq and CUT&RUN in day 6 RPCs. We performed ChIP-seq with (*Mettl3^KO^* rescued with HA-METTL3) and without Dox (*Mettl3^KO^*) in *Mettl3^KO^*;HA-METTL3^W^*^T^* Rx:GFP RPCs at day 6, using both HA antibody (Abcam, Ab9110), and a validated METTL3 antibody (Bethyl, A301-567A) respectively. Following integration of mapped peaks from HA and METTL3 ChIP-seq (+Dox), subtracted by non-specific peaks in *Mettl3^KO^* (-Dox), we identified 314 high-stringency METTL3-specific binding peaks, which we annotated to 222 genes (**Fig 2C**). We then integrated ChIP-seq with RNA-seq performed in parallel. We found no change in transcripts at METTL3 chromatin-bound genes in *Mettl3^KO^* (-Dox) and rescued *Mettl3^KO^* (+Dox) RPCs, indicating that METTL3 chromatin binding was non-functional in a transcriptional context in RPCs (**Fig S2A-C**). In addition to ChIP-seq, we also performed CUT&RUN with another validated METTL3 antibody (Abcam, Ab195352) in *Mettl3^KO^* (-Dox) and METTL3-rescued *Mettl3^KO^* (+Dox) RPCs. After exclusion of the non-specific binding peaks present in *Mettl3^KO^*, we identified 126 METTL3-specific binding peaks (**Fig 2D**). We did not observe correlation of H3K4me3 or H3K27me3 modifications at METTL3-specific chromatin targets, which indicates that METTL3 chromatin binding does not alter bivalent chromatin marks at corresponding sites (**Fig S2D**). Consistent with ChIP-seq, integration of CUT&RUN with RNA-seq (**Fig S2E**) did not show a correlation with METTL3-bound chromatin targets and DEGs between *Mettl3^KO^* (-Dox) and rescued *Mettl3^KO^* (+Dox) RPCs (**Fig 2E**). Taking together, we conclude that METTL3 chromatin binding in RPCs is uncommon, and that METTL3 chromatin targets are not functional: they are not associated with changes in bivalent histone modifications or transcript expression in RPCs.

### Genome-wide quantitative m6A mapping in mESC-derived RPCs reveals regulatory m6A modifications in eye field transcription factor RNA *Six3* at 3’UTR

Having established that METTL3 chromatin targets in RPCs are not associated with changes in bivalent chromatin marks or transcription, we sought to investigate how “canonical” METTL3-driven m6A modifications contribute to retinal development. To this end, we employed GLORI to obtain globally absolute-quantitative m6A maps at single-base resolution in day 4 WT and *Mettl3^KO^* organoids (performed in duplicate per WT and KO group) (Liu C 2023). After integrating WT and *Mettl3^KO^* duplicates separately, we found that WT and *Mettl3^KO^* organoids harbor 28,689 and 11,881 high-confidence m6A sites, respectively. Comparing these, we obtained 27,832 METTL3-dependent m6A sites (m6A level WT > *Mettl3^KO^*). GLORI-based mapping performed as expected, as sequence analysis of m6A sites lost in WT vs *Mettl3^KO^* revealed the presence of the conserved DRACH motif (**Fig 3A**). Among WT and *Mettl3^KO^* organoid transcripts (day 4, cultured in RPC differentiation conditions) centered on WT mRNAs with m6A marks, we saw a reduction in m6A levels (**Fig S3A**). Indeed, at these WT vs *Mettl3^KO^* m6A loss sites, m6A modification levels were low (10% < m6A level ≤ 40%) and medium (40% < m6A level ≤ 70%)(Liu, Sun et al. 2023) (**Fig S3A**). When analyzing all m6A modifications present in each WT and *Mettl3^KO^* organoid groups, we observed an overall loss in m6A level, as expected (**Fig 3B**). We analyzed distribution of m6A modifications across RNA domains, including enrichment at the 3’UTR (**Fig 3C**), consistent with the previous reports (Dominissini D 2012, Meyer KD 2012, Linder B 2015). Integration of GLORI and RNA-seq datasets from WT and *Mettl3^KO^* RPCs at day 4 revealed that m6A marks at 5’UTR, 3’UTR and CDS in *Mettl3*-dependent differentially regulated RNAs, which included EFTF transcripts such as *Lhx2*, *Six3* and *Rax*. *Lhx2* carries m6A modifications both at 5’UTR and 3’UTR, *Six3* with m6A at 3’UTR, and *Rax* at 5’UTR (**Fig S3B)**.

**Figure 3.**
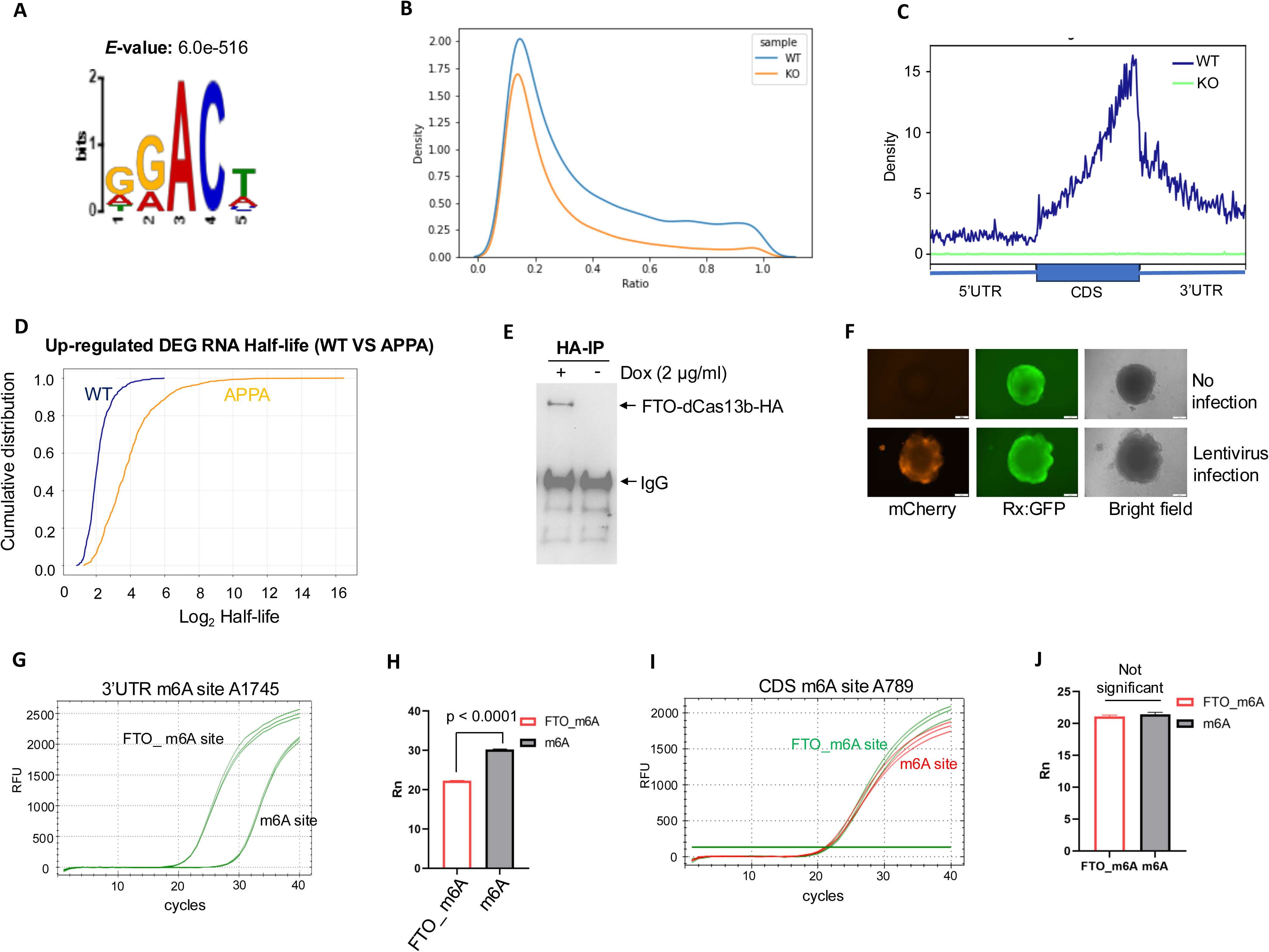
METTL3 regulates gene expression primarily through m6A-mediated RNA stability. (**A**) Motif analysis of lost m6A sites (n=27,832) identified by GLORI in WT vs. *Mettl3*^KO^ organoids after 4 days in RPC differentiation conditions reveals m6A-specific DRACH consensus motif, n=3 independent experiments. (**B**) Curve graph of m6A modification ratio in WT and *Mettl3*^KO^ organoids after 4 days in RPC differentiation conditions, n=3 independent experiments. (**C**) Metagene profiles showed the distribution of m6A sites between WT (n=28,689) and *Mettl3^KO^* (n=11,881) WT and *Mettl3*^KO^ organoids after 4 days in RPC differentiation conditions, n=3 independent experiments (**D**) Catalytic mutant *Mettl3* (APPA) enhances mRNA stability of up-regulated differentially expressed genes (DEGs) in *Mettl3*^KO^ organoids versus *Mettl3* WT organoids. RNA stability assay was performed via actinomycin D treatment at day 6 from *Mettl3*^KO^ organoids reconstituted with Flag-METTL3^WT^ or Flag-METTL3^APPA^ lines for 0, 2, 4, 6, 8 hr, and total RNA was subjected to mRNA-seq. (**E**) FTO-dCas13b-HA expression detected by HA immunoprecipitation followed by HA antibody detection in Rx:GFP mESCs with or without Dox (2μg/ml) treatment. (**F**) Fluorescence microscopy images show high infection efficiency in day 6 Rx:GFP^+^ RPCs when infected at day 0 with Lenti-mCherry gRNAs, n=3 independent experiments. (**G, H**) Single-base elongation and ligation-based qPCR amplification (SELECT) for the effect of FTO-dCas13b-mediated m6A removal at A1745 site of *Six3* 3’UTR increased *Six3* transcript level, n=3 independent experiments. In contrast, (**I,J**) SELECT assay showed that m6A removal at A789 site of *Six3* CDS had no effect on *Six3* mRNA level, n=3 independent experiments.

To interrogate the functional role of *Mettl3*-dependent m6A modifications in transcripts relevant to retinal development, we performed CRISPR-dCas13b mediated m6A engineering with FTO, an RNA m6A demethylase (Liu, Dou et al. 2020). We used gRNAs to guide FTO-dCas13b to remove m6A at *Six3* transcripts, followed by single-base elongation- and ligation-based qPCR amplification method (SELECT assay) to both detect m6A removal assess the effect of m6A removal on mRNA expression (Xiao, Wang et al. 2018). To this end, we stably expressed Dox-inducible HA-tagged FTO-dCas13b in Rx:GFP mESCs (**Fig 3E**). In parallel, we designed dCas13b gRNAs to target m6A sites present at CDS and 3’UTR of *Six3*, and non-target gRNA as control (**Table S1**). The gRNAs were cloned into a Lenti-guide-mCherry plasmid and the lentivirus containing gRNAs were infected into FTO-dCas13b Rx:GFP mESCs and mROs. More than 95% transduction efficacy was seen on day 6 Rx:GFP+ RPCs following day 0 (mESC) infection (**Fig 3F**). HA-tagged FTO-dCas13b was induced by Dox (2µg/ml) treatment in day 4 RPCs. SELECT showed that at A1745 (X3) site, gRNA-delivered FTO-dCas13b removed m6A, which led to a reduction in cycle number compared to the non-target gRNA control (**Fig 3G, H)**). qPCR CT with FTO-dCas13b m6A demethylation showed increased *Six3* mRNA level vs. non-target gRNA control. In addition, we targetedA1777 (X1 site) and A1798 (X2 site) sites, both in the 3’UTR, of *Six3* with FTO-dCas13b and found a similar pattern, with increased *Six3* mRNA expression vs control (**Fig S3C, D**). Finally, we targeted a non-3’UTR region, A789 (X4 site at CDS), and we did not see any change in *Six3* mRNA level (**Fig 3I, J**). Together, these results indicate m6A modifications in the *Six3* 3’UTR regulates its expression.

### METTL3 regulates gene expression primarily through m6A-mediated RNA stability

Following genome-wide quantitative m6A mapping and identification of METTL3-regulated DEGs (i.e. upregulated mRNAs in *Mettl3^KO^* organoids vs. WT organoids harboring Rx:GFP+ RPCs), we next asked if METTL3 m6A catalytic activity regulates RNA stability in METTL3-dependent transcripts. To this end, we performed RNA stability assays. Specifically, we cultured Flag-METTL3 WT or Flag-METTL3 APPA (catalytic mutant)-expressing *Mettl3^KO^* mESCs for 6 days in mESC-to-RPE differentiation conditions. After 6 days, we performed an RNA stability assay with Actinomycin D treatment at 0, 2, 4, 6, 8h timepoints and total RNA was subjected to mRNA-seq. The results showed that the expression of METTL3 APPA mutant lengthens half-life of METTL3-regulated mRNAs (**Fig 3D**). Taken together, these data suggest that METT3-mediated m6A modification of RNAs may be the primary mechanism whereby METTL3 influences mESC-to-RPC differentiation.

### Acute METTL3 inactivation in RPCs reveals a regulatory METTL3-*Ythdf1* protein-RNA interaction and leads to impaired retinal differentiation

We next investigated METTL3-bound RNAs that harbor m6A modifications in RPCs, as an initial step for integrative analysis to identify METTL3 RNA targets that regulate retinal development. We performed eCLIP using a validated METTL3 antibody (Abcam, Ab195352) with 1x10^8^ of mESC-derived WT (Rx:GFP+) or *Mettl3^KO^* (Rx:GFP-) organoids on day 6 (Van Nostrand, Pratt et al. 2016). A total of 2584 high stringency METTL3 target RNAs were identified by excluding peaks from *Mettl3^KO^* organoids (**Fig 4A**). We identified histone clusters which served as a quality control for eCLIP assay, which have been previously reported target RNAs for METTL3 (**Fig S4A**) (Van Nostrand EL 2016). GO term pathway analysis of the 2584 METTL3-bound RNAs showed that the most enriched pathway is RNA binding (p = 2.60e^-44^) and followed by ribonucleoprotein complex and mRNA processing (**Fig 4B**).

**Figure 4.**
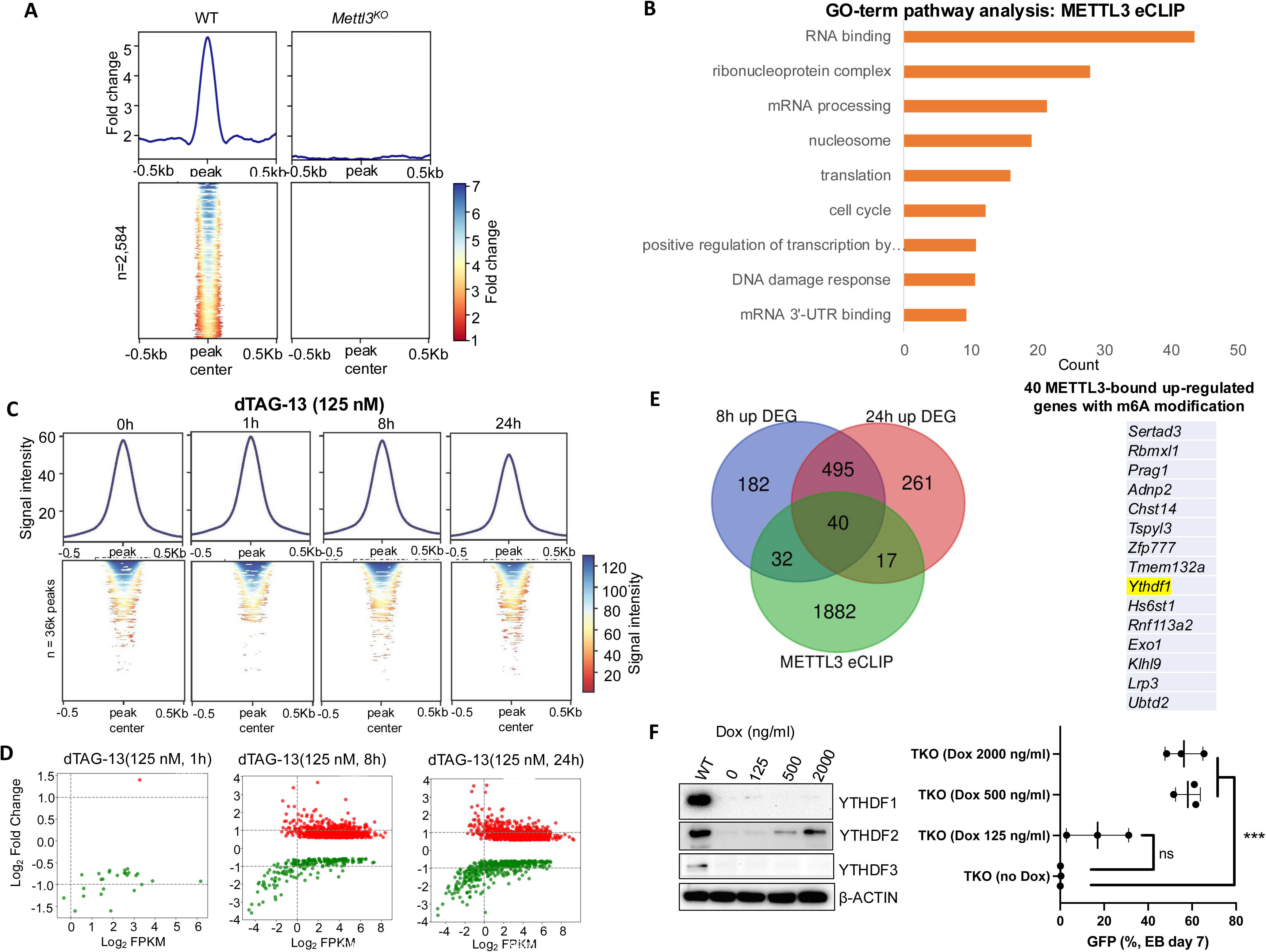
Targeted acute METTL3 loss in RPCs leads to impaired retinal development *in vitro* and *in vivo*. **(A)** Heatmaps and metagene profiles showing METTL3-eCLIP enrichment of WT-specific METTL3-binding peaks were reduced in *Mettl3^KO^* organoids (negative control) after 6 days under RPC differentiation conditions. X-axis shows distance from center of peak in base pairs. Heatmaps are ranked according to METTL3 enrichment in WT RPCs in descending order, n=3 independent experiments. (**B**) GO-term analysis of METTL3 direct target RNAs by METTL3-eCLIP. (**C**) ATAC-seq heatmaps following acute METTL3 degradation by dTAG-13 (125nM) treatment for 1hr, 8hr and 24hr, no treatment (0hr) as control, n=3 independent experiments. (**D**) Volcano plot showing up-regulated DEGs (red dots) and down-regulated DEGs (green dots) upon acute degradation of METTL3 by dTAG13 (125nM) after 1hr, 8hr and 24hr treatment, n=3 independent experiments. (**E**) Integrative analysis showed 40 RNAs specifically bound by METTL3 (eCLIP) that were significantly up-regulated (RNA-seq) by acute degradation of METTL3 at both 8hr and 24hr treatment. (**F**) *Ythdf1&2&3* triple-knockout (TKO) RGSC mESC-derived RPCs phenocopied *Mettl3^KO^* RPCs. *Ythdf1&2&3* TKO mESC lines harboring Dox-inducible YTHDF2 was generated in dual reporter Rx:GFP;Sox1:mCherry RGSC mESCs and dose-dependent assay was performed and detected by WB with YTHDF1, YTHDF2 and YTHDF3 antibodies, β-ACTIN as loading control (left panel). Flow cytometry of Rx:GFP expression of RPCs (day 6) of the TKO line treated with different dosages of Dox (right panel, Dox inducible YTHDF2 rescue assay). Paired group student t test was performed. Ns, no significance, n=3 independent experiments.

In our Dox-inducible rescue complementation system, we observe mESC-to-RPC differentiation defects after conditional *Mettl3* inactivation for 4-6 days (**Fig 1B**). However, the acute consequences of METTL3 depletion in RPCs remain unknown, especially with respect to chromatin accessibility and direct METTL3 RNA targets we identified in **Fig 4A**. To address these knowledge gaps, we utilized a degron-based targeted protein degradation system (Nabet, Roberts et al. 2018). FKBP12^F36V^-HA was knocked into the C-terminal end of the two endogenous *Mettl3* alleles in Rx:GFP mESCs. Acute degradation of endogenous METTL3 was induced by dTAG-13 treatment. On day 6 of RPC differentiation, both dose- and time-dependent, near complete degradation of FKBP12^F36V^-HA-METTL3 as quickly as 2h post dTAG-13, or as low as 62.5nM of dTAG-13 (**Fig S4B**). For downstream studies, we chose 125nM dTAG-13 as the dose for time dependent assay: as short as 4hrs of treatment, complete METTL3 degradation was observed (**Fig S4C**).

Since METTL3-dependent m6A modifications have been implicated in regulation of chromatin accessibility (Deng, Zhang et al. 2022), we explored whether acute METTL3 degradation alters chromatin accessibility in RPCs. We used ATAC-seq and RNA-seq in FKBP12^F36V^-HA knock in Rx:GFP mESC-derived RPCs at 1hr, 8hr and 24hr dTAG-13 (125nM) treatment, no treatment (0hr) as control. Interestingly, while we did not observe a change in chromatin accessibility at 8h, and 24h following acute METTL3 degradation (**Fig 4C**), we saw transcriptional changes at each of these timepoints by RNA-seq (**Fig 4D**). Thus, chromatin accessibility is uncoupled from rapid transcriptional changes that follow acute METTL3 loss.

To further explore the significance of rapid transcriptional changes from acute METTL3 degradation, we analyzed DEGs from RNA-seq after dTAG-13 treatment (**Fig 4D**). We saw significantly more DEGs at 8h and 24h after dTAG-13 than those at 1h (**Fig S4D**). Classically, m6A modifications are thought reduce RNA stability; thus loss of m6A leads to increased expression of corresponding RNA (Wang, Lu et al. 2014). In this vein, we integrated RNA-seq, GLORI and METTL3-eCLIP and identified 40 RNAs that were m6A-methylated, directly bound by METTL3, and significantly up-regulated by acute degradation of METTL3 at both 8hr and 24hr treatment (**Fig 4E**). Interestingly, we found that *Ythdf1* RNA, which encodes an m6A reader, is bound by METTL3, rapidly upregulated with METTL3 degradation, and harbors m6A modifications (**Fig S4E, F**). By GLORI, we found that *Ytdhf1* m6A modifications decreased after 8hr dTAG-13 treatment (**Fig S4F**). In addition, we found *Ythdf2* RNA was also upregulated by acute METTL3 loss, but it was not bound by METTL3 by eCLIP (**Fig S4E**). This finding suggests a potential METTL3-*Ythdf1* protein-RNA feedback loop in RPCs. The YTHDF family consists of YTHDF1, 2, 3 and these three m6A readers have overlapping roles in mESCs (Lasman, Krupalnik et al. 2020). With this in mind, we generated a triple *Ythdf1/2/3* knockout (TKO) in Rx:GFP mESCs and differentiated them to RPCs. TKO mESCs phenocopied *Mettl3^KO^* mESCs in their inability to generate Rx:GFP+ RPCs.

Importantly the TKO mESC-to-RPC defect was rescued with Dox-induced YTHDF2 expression (**Fig 4F**). Together, our findings show that during acute METTL3 degradation in RPCs, chromatin accessibility is uncoupled from transcriptional changes, but changes in the m6A deposition on METTL3 target RNAs occur rapidly. Indeed, a METTL3 target RNA, *Ythdf1*, is rapidly regulated at the m6A and transcription levels following METTL3 loss. Finally knockout of the entire *Ythdf* family (that encode three m6A proteins with redundant mESC function) phenocopies *Mettl3* loss to impair mESC-to-RPC differentiation, suggesting a cell fate function for the METTL3-*Ythdf1* protein-RNA interaction.

## Discussion

How epitranscriptome controls differentiation is a central question in development. Here, we show that METTL3, the main m6A writer, regulates the mESC-to-RPC and RPC-to-retina transcriptional programs through mechanisms independent of chromatin accessibility and its ability to directly engage chromatin. Interestingly, we found that in mESC-to-RPC organoid retinal differentiation conditions, *Mettl3* loss did not delay pluripotency exit, as reported by others (Jin, Zuo et al. 2021), but instead found a lineage-specific effect whereby *Mettl3^KO^* mESCs differentiated to cells with pan-neuroectoderm markers such as *Sox2*, *Otx2*, and *Pax6*; but showed reduced acquisition of eye field/retinal markers such as *Rx*, *Lhx2*, and *Six3*.

METTL3 is observed predominantly in nuclear speckles, although m6A methyltransferase activity has been detected in both nuclear and cytoplasmic fractions, including in RPCs.(Lin S 2016) A functional nuclear localization signal (NLS) on METTL3 has been reported and NLS mutant of METTL3 retains in cytoplasm (Schöller E 2018). To further confirm the cytosolic METTL3 function in RPC differentiation, we reconstituted NLS mutant METTL3 (AKKSRK to AGGSGG or ARRSRR) in *Mettl3^KO^* mESCs and confirmed its expression by Western blotting. However, the NLS mutant METTL3 could not rescue defective mESC-to-Rx:GFP+ RPC differentiation of *Mettl3^KO^* organoids (**Fig S3E, F**). This finding supports our conclusion that nuclear METTL3 is required for mESC-to-RPC differentiation.

Recent reports have described a diversity of non-canonical epigenome-epitranscriptome crosstalk of METTL3, including its binding to the promoters of heat shock gene loci (Knuckles P 2017), senescence associated secretory phenotype (SASP) genes (Liu P 2021), CAATT-box binding motif containing genes (Barbieri I 2017) and the intracisternal A Particle (IAP) type family (Xu W 2021), each with distinct contextual functions. In contrast, we found *Mettl3* plays a potent role in retinal differentiation through the epitranscriptome, but with limited crosstalk to the epigenome observed.

First, while we saw that METTL3 bound to a modest number of chromatin sites, we did not see corresponding transcriptional changes at those sites, indicating that METTL3-to-chromatin binding may not be functional in mESC-to-RPC fate choice. In addition, our degron-based strategy uncovered, for the first time, events immediately following METTL3 degradation in RPCs on a timescale of hours. While we saw changes in transcription and m6A modifications, we did not observe major alterations to chromatin accessibility, indicating an uncoupling between the epigenome and epitranscriptome in this context.

Our seminal dCas13b-FTO m6A engineering in retinal cells reveals the function of 3’UTR m6A modifications in the EFTF transcript, *Six3*. We saw that the removal of m6A marks at 3’UTR of *Six3* enhances its mRNA stability. m6A modifications in the 3’UTR are recognized by YTHDF2 to regulate mRNA degradation (Wang X 2014), while synergistic interactions between YTHDF1 and YTHDF3 allow recognition of m6A in the 3’UTR to facilitate translation (Wang X 2015, Shi H 2017). YTHDF1/2/3 are redundant in their roles to promote degradation of m6A modified RNAs (Zaccara S 2020, Niu, Che et al. 2022, Zaccara S 2024). Interestingly, m6A depletion via 3’UTR mutations of *insulin* mRNA or indirectly by mutating *Mettl3* decreased insulin protein production, leading to aberrant homeostasis and diabetic-like phenotype (Wilinski 2023). Furthermore, through integrative analyses of METTL3 eCLIP, GLORI and identification of rapidly altered transcripts following acute METTL3 degradation in RPCs, we discovered a regulatory protein-RNA interaction between METTL3 and *Ythdf1*.

Our studies include limitations that would point to areas for future investigation. While we report a METTL3-*Ythdf1* protein-RNA interaction on day 6 RPCs for the first time, further investigation of this interaction during mESC-to-RPC differentiation would provide further insights. Through GLORI, SELECT, dCas13-FTO m6A engineering, it will be important to identify functions for METTL3-direct m6A catalytic targets on RNAs whose METTL3-dependent destabilization leads to loss of retinal differentiation in *Mettl3^KO^* mESCs. Further, a deeper understanding of how rapid transcriptional changes occur following acute METTL3 degradation in the absence of major changes in chromatin accessibility is important.

While we did not find many METTL3 binding sites on chromatin, and that METTL3 chromatin targets did not show differential transcription, we cannot rule out an indirect effect of METTL3 on chromatin modification, given that we observed METTL3-dependent changes in histone modifications. Indeed, several reports identified specific crosstalk between m6A RNA modifications and histone modifications, including H3K36me3 (Huang 2019), H3K9me2 (Li Y 2020), H3K9me3 (Liu 2021) and K3K27Ac (Huang, Zhang et al. 2025). A recent report demonstrated crosstalk between m6A RNA methylation and 5mC DNA methylation by METTL14-mediated recruitment of DNMT1 to chromatin and the balance between these two marks controls key differentiation genes in mESCs (Quarto, Li Greci et al. 2025). Potential METTL3-associated chromatin modifier proteins could be explored in the future. Do m6A-modified RNAs show association with chromatin sites? Some m6A-modified chromatin-associated RNAs (caRNAs) can bind directly to DNA to regulate chromatin state and downstream transcription (Liu, Dou et al. 2020). Future studies that combine GLORI, dCas13b-FTO m6A engineering, CRISPR-Cas9 DNA editing at RNA-chromatin contacts, along with a suite of RNA-chromatin profiling technologies could shed further light on how or whether particular m6A-modified RNAs functionally associate with chromatin and influence lineage specification (Khlebnikov, Nikolskaya et al. 2025).

Together, our studies provide seminal advances in our understanding of the essential function of METTL3 and the epitranscriptome during tissue development: distinct from its canonical m6A regulation, METTL3 targets on chromatin and epigenome regulation are unexpectedly uncoupled from transcription.

## Materials and Methods

### mESC maintenance, induced retinal neuroectoderm differentiation and plasmid electroporation

Rx:GFP ESC maintenance, induced retinal neuroectoderm differentiation and plasmid electroporation were performed as described previously (Eiraku M 2011, Li, Mao et al. 2020, Li, Huang et al. 2021). WT or mutant *Mettl3* inserts (APPA, W475A and N477A) were subcloned to doxycycline inducible or constitutive PiggyBac plasmids as described previously (Xu, Li et al. 2021). For Dexamethasone inducible WT or APPA Mettl3 plasmid, glucocorticoid receptor ligand binding domain (GRBD) insert from pPyCAG-cGR-IP plasmid (RIKEN, #RDB10440) was subcloned to constitutive PiggyBac plasmid to generate GRBD-FLAG-METTL3^WT^ or GRBD-FLAG-METTL3^APPA^ fusion protein (Murakami, Gunesdogan et al. 2016). For CRISPR/Cas9 mediated *Mettl3* knockout, two independent *Mettl3* guide RNAs (GTGCTTATTGATAATTCGTCTGG and TGTGAAGCTGGTTAAAGCTCAGG) were used to generate independent *Mettl3* KO ES clones.

#### ChIP-seq

Chromatin immunoprecipitation followed by sequencing (ChIP-seq) was performed using a slightly modified version of ChIP Assay Kit (Millipore) based protocol as described previously (Li, Mao et al. 2020, Li, Huang et al. 2021). Day 6 *Mettl3*^KO^ mROs with or without Dox (2 µg/ml) treatment were trypsinized into single cells and then total cell number 2×10^7^ cells for each ChIP were fixed with 10% paraformaldehyde (Sigma) at room temperature for 10 min and quenched the reaction using glycine at final concentration of 0.125 M. The cell pellets were further sonicated to 300–500 bp in SDS lysis buffer by using a Qsonica Biorupter. 15 μg ChIP grade anti-HA antibody (Abcam, ab9110) and anti-METTL3 (Bethyl, A301-567A) were used for ChIP assay. The ChIP-DNA were sent to University of Michigan DNA Sequencing Core for quality control, library preparation, and sequenced using Illumina NovaSeq S4 Libraries platform.

### CUT&RUN

Day 4 WT and *Mettl3*^KO^ mROs were harvested and trypsinized to single cell suspension. 2×10^6^ cells were subjected to CUT&RUN per antibody as previously described (Li, Mao et al. 2020, Li Q 2020, Li, Huang et al. 2021). A CUT&RUN kit (EpiCypher, 14-1048) was used as per manufacture’s protocol and the following antibodies were used: METTL3 antibody (Abcam, Ab195352), H3K4me3 antibody (Millipore, 07-473) and H3K27me3 antibody (ActiveMotif, 39155).

### GLORI for absolute quantification of transcriptome-wide m6A at single-base resolution

GLORI was performed as reported previously (Liu C 2023), slightly modified, as follows: 100 µg of total RNA from WT or *Mettl3*^KO^ mROs (day 4) passed QC (< 200nt of RNA) using MEGA clear transcription clean-up kit (Ambion, AM1908), and then underwent PolyA+ selection, fragmentation, glyoxal protection, deamination and deprotection. Final RNA was purified by ethanol precipitation and twice clean-up by RNA Clean & Concentrator kit. The RNAs were sent to University of Michigan DNA Sequencing Core for direct RT and library construction, and sequencing using Illumina NovaSeq S4 Libraries platform with PE read lengths of 2 ×150 base pairs.

### m6A Engineering by FTO-dCas13b and Readout by SELECT

To investigate m6A modification function at mRNAs such as *Six3* 3’UTR, we first generated a Dox-inducible, HA-tagged FTO-dCas13b (FTO from Addgene 134781 and dCas13b from Addgene 103865) stable overexpression line in Rx:GFP mESCs. HA-FTO-dCas13b expression level was determined by HA-IP and western blot by HA antibody (Abcam ab9110) upon Dox (2µg/ml) treatment. In parallel, design dCas13b gRNAs were used to target all m6A sites at the 3’UTR of *Six3*, and non-target gRNA as control. The gRNAs were cloned into Lenti-guide-mCherry plasmid (Addgene 170510). gRNAs used in this m6A removal assay are listed in Supplementary table 1.

Optimization of infection efficacy of the lentivirus containing gRNAs were determined by time course assay (day 0, day 2 and day 4) of Rx:GFP RPCs and viewed at day 6 by mCherry and GFP under fluorescence microscopy. The detection of m6A at targeted sites was based on SELECT method previously developed by Xiao et al (Xiao Y 2018). In all, 1.5 µg total RNA was first mixed with 100 nM Up primer, 100 nM Down primer, and 5 µM dNTP in 17 µL 1v CutSmart buffer. The mixture was then annealed in the following programs: 90 °C (1 min), 80 °C (1 min), 70 °C (1 min), 60 °C (1 min), 50 °C (1 min), and 40 °C (6 min). Next, 17 µL annealing products were incubated with 3 µL enzyme mixture containing 0.01 U Bst 2.0 DNA polymerases, 0.5 U SplintR liagase, and 10 nmol ATP. Subsequently, the final mixture (totally 20 µL) above was incubated at 40 °C for 20 min, denature at 80 °C for 20 min. SELECT qPCR was carried out using IQ SYBR Green Supermix (BioRad, 170-8882AP). Primers used in SELECT measurement are listed in **Supplementary Table 1**.

### METTL3 eCLIP

METTL3 eCLIP was performed as described previously (Van Nostrand EL 2016). In brief, UV crosslinked WT and Mettl3 KO Rx:GFP RPCs were collected and resuspended in 600µl of RIPA lysis buffer with freshly added Protease Inhibitor Cocktail (Roche, 11873580001) and 1 μl of Superase-In RNase inhibitor (Thermo Fisher, AM2694)) for 15 min on ice and then sonicated with Qsonica 800R (30 s, on; 30 s off; 20 minutes, 30% amplitude). Sonicated lysates were centrifuged at 13,000 × g for 15 min at 4 °C. Clarified lysates were incubated with 15µl METTL3 antibody (Abcam, Ab195352) at 4 °C overnight. Two percent of lysate was saved as size-matched input. 30µl of Protein A Dynabeads (Thermo Fisher, 10001D) were added into IP tubes and incubated at 4 °C for 2 hrs. The immunoprecipitated (IP) samples were washed with low salt buffer (20 mM Tris-HCl pH 8, 150mM NaCl, 2 mM EDTA, 1% Triton X-100, 0.1% SDS, freshly added RNase Inhibitor at 1:1000 dilution) twice and then washed with LiCl wash buffer (10 mM Tris-HCl pH 8, 0.25 M LiCl, 1 mM EDTA, 1% NP-40, 0.25% sodium deoxycholate, freshly added RNase Inhibitor at 1:1000 dilution) twice. The IP samples underwent final washes twice with TE buffer (10 mM Tris-HCl pH 80 and 1mM EDTA, freshly added RNase Inhibitor at 1:1000 dilution). IP samples were eluted from beads with 300 µl elution buffer (50 mM Tris-HCl pH 8, 150mM NaCl, 1 mM EDTA, 1% SDS, 50mM NaHCO3 and freshly added RNase Inhibitor at 1:1000 dilution and 7µl of Proteinase K) at 37 °C for 30 minutes. 500µl Trizol was added to extract RNA and then digested with DNase (Thermo Fisher, NC1424104) at 37 °C for 30 minutes. The Isopropanol precipitated RNA was resuspended in 17.5 µl of RNase/DNase free water at 55 °C for 5 minutes. Libraries were amplified using standard RNA library prep kit by UM AGC. The quantity and quality of the final libraries were assessed using a Bioanalyzer (Agilent Technology Inc). All samples were sequenced by PE150 on the Illumina NovaSeq shared flow cell sequencer. Two biological replicates were conducted for each experiment.

### Targeted METTL3 Degradation by dTAG-13

The dTAG system (Nabet B 2018) was adapted to induce FKBP12^F36V^-METTL3 acute degradation. First, we constructed a donor plasmid of FKBP12^F36V^ -2XHA into Dox-inducible PiggyBac plasmid with 250bp of left and right homology arm right before and after stop codon of *Mettl3*. *Mettl3* gRNA was constructed into PX459 plasmid carrying a puromycin selection marker. Then, the donor plasmid was co-transfected with PX459-gRNA plasmid into Rx:GFP mESCs by Nucleofector and 48 post transfection, the cells were selected with puromycin 1.5µg/ml for 5 days. The knock-in stable colonies were screened by genotyping and western blot with both HA antibody and METTL3 antibody. Both the dose-dependent assay and time course assay of dTAG-13 treatment using the knock-in Rx:GFP mESC cell line were performed to determine the ideal dose and time point of dTAG-13 treatment to achieve acute METTL3 degradation. From the above results, we used dTAG-13 125nM to treat differentiated Rx:GFP mESC-derived RPCs at day 6 for 0h, 1h, 8h and 24h and performed ATAC-seq and paired mRNA-seq.

## Resource availability

Lead contact

Further information and requests for resources and reagents can be directed to the lead contact, Rajesh C. Rao.

Materials availability

All unique reagents generated in this study are available from the <u>lead contact</u>.

Data and code availability

GEO Accession Numbers associated with this manuscript:

GSE287489 for XUY9572: Nanopore seq

GSE287484 for 7120-JX: mRNA-seq

GSE287232 for 3912-JX: mRNA-seq

GSE287253 for 11904-JX: mRNA-seq

GSE287148 for 9022-JX: CUT&RUN

GSE287482 for 3258-JX: ChIP-seq

GSE287485 for 9446-JX: m6A GLORI

GSE287488 for 12204-JX: m6A GLORI

GSE287302 for 11646-JX: METTL3 eCLIP

GSE287487 for 11870-QL: ATAC-seq

## Acknowledgments

R.C.R. was supported by the National Eye Institute (NEI, R01EY030989), Research to Prevent Blindness (RPB) Career Advancement and Unrestricted Departmental Awards, A. Alfred Taubman Medical Research Institute, the Beatrice and Reymont Paul Foundation, March Hoops to Beat Blindness, and Leonard G. Miller Endowed Professorship and Ophthalmic Research Fund at the Kellogg Eye Center. Additional support for this research was provided by Grossman, Elaine Sandman, Marek and Maria Spatz (endowed fund), Greenspon, Dunn, Avers, Boustikakis, Sweiden, and Terauchi research funds. The University of Michigan Vision Research Center (P30EY007003) provided some Core Services. J.L. was supported by NIGMS R35HG011279. We thank Charukesi D. Sivakumar, BS for her assistance on the Graphical Abstract (generated with Biorender).

## Author Contributions

Conceptualization, R.C.R.; Methodology, X.J., Y.H.,Z.H., Q.L., J.L., and R.C.R.; Software and Computational Analysis, Y.H. and Z.H.; Investigation, X.J., Y.H.,Z.H., Q.L., J.L., and R.C.R; Resources, J.L., and R.C.R.; Writing – Original Draft, X.J., Q.L. and R.C.R.; Writing – Review & Editing, X.J., Q.L., J.L., and R.C.R.; Supervision, R.C.R.; Funding Acquisition, R.C.R.

## Declaration of Interests

All other authors have no competing interests to declare.

## Supplemental Materials and Methods

### RNA Isolation, Reverse Transcription, and Quantitative Real-Time PCR (RT-qPCR)

Trizol reagent (ThermoFisher Scientific) and RNeasy mini kit (QIAGEN) were used for RNA isolation from ESCs and mROs. 1.0 μg total RNA was reverse transcribed to cDNA using high-capacity RNA-to-cDNA kit (ThermoFisher Scientific). RT-qPCR was performed using Taqman probes (ThermoFisher Scientific). Gapdh, β-actin was used as internal control for normalization. Probe sequences are available on request. Data were automatically analyzed using Bio-Rad CFX manager software.

### Whole-Cell Lysate Preparation, Western Blotting and Cell Fractionation Assay

ESCs or mROs were lysed with RIPA buffer (Pierce) in the presence of EDTA-free protease inhibitor cocktail (Roche). Protein concentration was determined using Pierce BCA protein assay kit (ThermoFisher Scientific). 50 μg of whole cell lysate was resolved on 4–20% precast gel (Bio-Rad) was transferred to 0.45 μm PVDF membrane (Millipore). The following primary antibodies were used for probing: anti-HA (1:10,000, Abcam, ab9110), anti-METTL3 (1:5000, Bethyl), anti-METTL3 (1:5,000, Abcam), anti-Flag (1:2,500, Sigma), anti-β-Actin-HRP conjugate (1:10,000, Abcam), anti-VDAC (1:2,500, Cell Signaling) and anti-LSD1 (1:2,500, Abcam). A nuclear/cytosol fractionation kit (ThermoFisher Scientific, cat#78833) was used to separate cytoplasmic and nuclear cell fractions in sorted day 6 Rx:GFP+ RPC cell population. Trypsinized cells were washed in PBS and re-suspended in cell lysis buffer, then treated according to the manufacturer’s protocol.

### Flow Cytometry

mROs (organoids) were combined and dissociated to single cell suspension by 0.25% trypsin-EDTA (Invitrogen). DMEM media with 10% FBS (Sigma, v/v) was used to inactivate trypsin, and cells were subjected to spin down at 300 g for 5 min. Cell pellet was washed and re-suspended in D-PBS (Invitrogen). Cells were immediately transferred to flow cytometer for data analysis (Sony SH800) or sorting (Bigfoot) for Rx:GFP+ RPCs. The dead cells in the cell population were gated out and percentage of Rx:GFP+ RPCs cells were calculated on live cells.

### Nanopore Sequencing

One microgram of poly(A)-selected RNA of Rx:GFP mRO (day 6) with replicates were subjected to the Nanopore direct RNA sequencing using kit (SQK-RNA002). Sequencing was carried out on a GridION Nanopore FLO-MIN106D flow cell for each library for ∼48h. Data was base-called with Guppy v4.0.11 base callers. Total reads (in millions) were 1.3 and 0.81 respectively. Detected m6A modification loci with MINES (m6A Identification using Nanopore Sequencing) uses a compilation of the four random models, each corresponding to a DRACH motif, AGACT, GGACA, GGACC, and GGACT. MINES uses Tombo’s fraction-modified values and coverage files as inputs and outputs a bed file of predicted sites. For more information, visit https://github.com/YeoLab/MINES.git.

### Bulk RNA-Seq

RNA extraction for RNA-seq was performed via combination of Trizol reagent (ThermoFisher Scientific) and RNeasy mini kit (QIAGEN). Total RNA extracted from day 0 of WT, *Mettl3*^KO^ and *Mettl3*^KO^+METTL3^WT-GRBD^ mESCs treated with or without Dex (10µM), Day 2, 4 and 6 of WT, *Mettl3*^KO^ and *Mettl3*^KO^+METTL3^WT-GRBD^ mROs treated with or without Dex (10 µM) or Day 6 of *Mettl3*^KO^ mROs treated with or without Dox (2 µg/ml) were sent to University of Michigan DNA Sequencing Core for RNA quality analysis, PolyA+ mRNA selection and library construction, and sequencing using Illumina NovaSeq S4 Libraries platform. Duplicate samples for each group were subjected to two rounds of independent library preparation and sequencing to avoid sample and batch effect of mRNA-seq.

### Quantification and Statistical Analysis

All experiments were independently repeated at least for 2 to 4 times with similar results, and data from one representative experiment are presented unless otherwise stated. Two-tailed unpaired Student’s *t* test was applied using GraphPad Prism (version 7.00) to determine whether the observed differences were statistically significant. Changes were considered statistically significant if *p* value was less than 0.05.

### Bioinformatics Analysis

All RNA-seq, ChIP-seq CUT&RUN, and ATAC-seq data analysis were according to previously described (Li et al. 2020; Li et al. 2021).

### GLORI m6A modification calling and annotation

R2 reads were used for m6a detection and quantification. BCL Convert Conversion Software v4.0 (Illumina) was used to generate de-multiplexed Fastq files for the Illumina NovaSeq library. Adapters were removed using trim_galore v.0.6.6 with parameters ‘-q 20 --stringency 1 -e 0.3 --length 35’. Seqkit v.0.13.2 was used to deduplicate PCR according to the 10bp UMIs of the reads with parameters ‘rmdup -s’. FASTX-Tool kit v.0.0.13 was used to remove the 10bp UMIs in the deduplication reads with parameters ‘fastx_trimmer -f 11’. GLORI-tools v1.0.0 was used to call m6a modifications from the processed reads with parameters ‘--combine --rvs_fac’.M6a modifications were annotated to the reference genome mm10 using the annotate PeakInBatch function in R package Modifications with introns were specifically identified using the intronsByTranscript function in R package ensembldb. Differential m6a modifications were identified using R package methylSig.

### eCLIP bioinformatic analysis

All eCLIP-seq data were processed using the CLIP Tool Kit (CTK) following a structured pipeline tailored for accurate analysis of eCLIP experiments. Initially, raw reads were preprocessed to remove low-quality sequences and adapter contamination using fastq_quality_filter from the FASTX-Toolkit with parameters -q 20 -p 80 and cut adapt with parameters -a AGATCGGAAGAGCACACGTCTGAACTCCAGTCAC -m 18. Preprocessed reads were then mapped to the mm10 mouse genome using Bowtie with parameters -q -phred33 -l 20 -n 2 -k 1 -m 1 --best --strata, ensuring high stringency and accuracy. Post-mapping, unique reads were retained and PCR duplicates were removed using the <u>tag2collapse.pl</u> tool, which collapses reads based on unique start positions. Crosslinking-induced truncations were identified using parseAlignment.pl with --mapq 30 for filtering high-quality alignments, followed by the generation of crosslink-centered clusters using <u>tag2cluster.pl</u>. Peak calling was performed with <u>tag2peak.pl</u> using the input as a control for robust identification of true binding sites. Finally, visualization-ready bigwig files were created from BAM files using the <u>bed2wig.pl</u> script with parameters --bw --ext 200 --rpm, enabling data exploration in tools like Integrative Genomics Viewer (IGV). This comprehensive pipeline ensures the reproducibility and high-resolution analysis of eCLIP data, enabling insights into RNA-binding protein interactions.

## Supplemental Figures and Figure Legends

**Figure S1, related to Figure 1:**
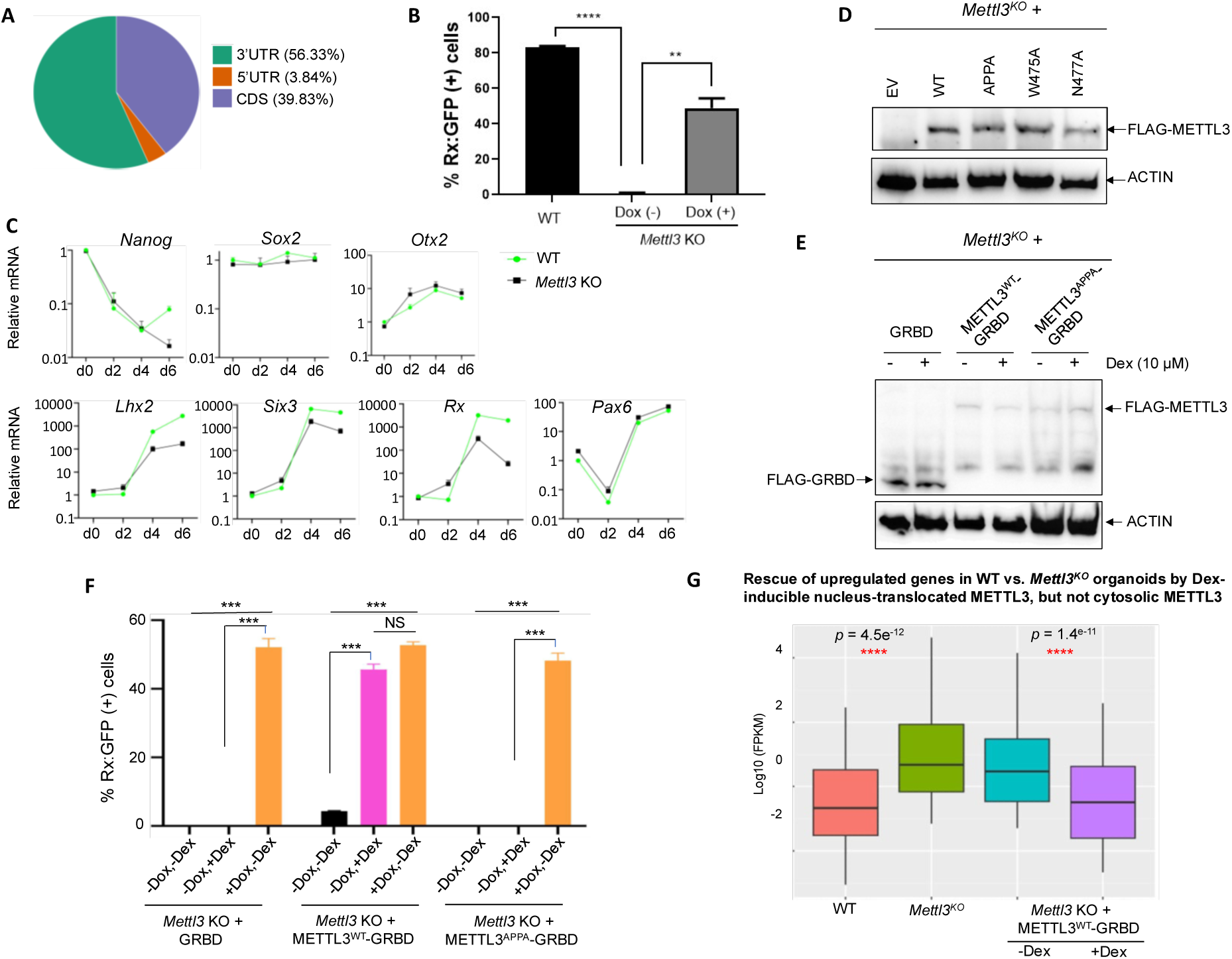
Nuclear METTL3 with intact catalytic activity is required for the mESC-to-RPC cell fate transition. **(A)** Genome distribution of m6A modifications in transcripts of day 6 Rx:GFP+ RPCs, detected via nanopore sequencing. PolyA-selected mRNAs from RPCs were subjected to nanopore sequencing for mapping m6A sites, n=2 independent experiments. (**B**) Quantification of Rx:GFP expression by flow cytometry (n=3 independent experiments). Rescue complementation *Mettl3*^KO^ mESCs with Dox-inducible METTL3^WT^ were differentiated to RPCs over 6 days. RPCs treated with and without Dox were analyzed by flow cytometry for Rx:GFP expression. **** P<0.0001, ** P<0.01, WT Rx:GFP as control. (**C**) RT-qPCR validation (n=3) of the RNA-seq data presented in Fig. 1C. (**D**) Reintroduction of the FLAG-tagged constitutive WT and catalytic mutants (METTL3^APPA^, METTL3^W475A^, METTL3^N477A^) METTL3 plasmids into *Mettl3^KO^* mESCs. EV: empty vector control. Western blot detected by FLAG antibody and ACTIN as loading control, n=3 independent experiments. (**E**) Constitutive expressions of FLAG-tagged GRBD, METTL3^WT^-GRBD and METTL3^APPA^-GRBD into *Mettl3^KO^* mESCs. Western blot detected by FLAG antibody and ACTIN as loading control, n=3 independent experiments. (**F**) Flow quantification of Rx:GFP expression at day 6 (n=3) under RPC differentiation conditions of *Mettl3^KO^* mESCs with constitutive expression of Flag-tagged GRBD, METTL3^WT^-GRBD and METTL3^APPA^-GRBD treated with or without Dox and Dex. +Dox induces METTL3^WT^ expression and +Dex induces translocation of cytosolic METTL3-GRBD to the nucleus. NS, no significance, n=3 independent experiments. (**G**) RNA-seq showed upregulated genes in WT or *Mettl3^KO^* cells (day 4) expressing empty vector (*Mettl3^KO^*), cytosolic METTL3-GRBD (-Dex), or nuclear METTL3-GRBD (+Dex). Upregulated genes in WT vs *Mettl3*^KO^ organoids and Dex-rescue (-Dex [cytosolic METTL3-GRBD] vs. +Dex [nuclear METTL3-GRBD]) (day 4) were compared by pairwise student t-tests (asterisks denote p < 0.05). All other pairwise comparisons were non-significant (p > 0.05), n=3 independent experiments.

**Figure S2, related to Figure 2:**
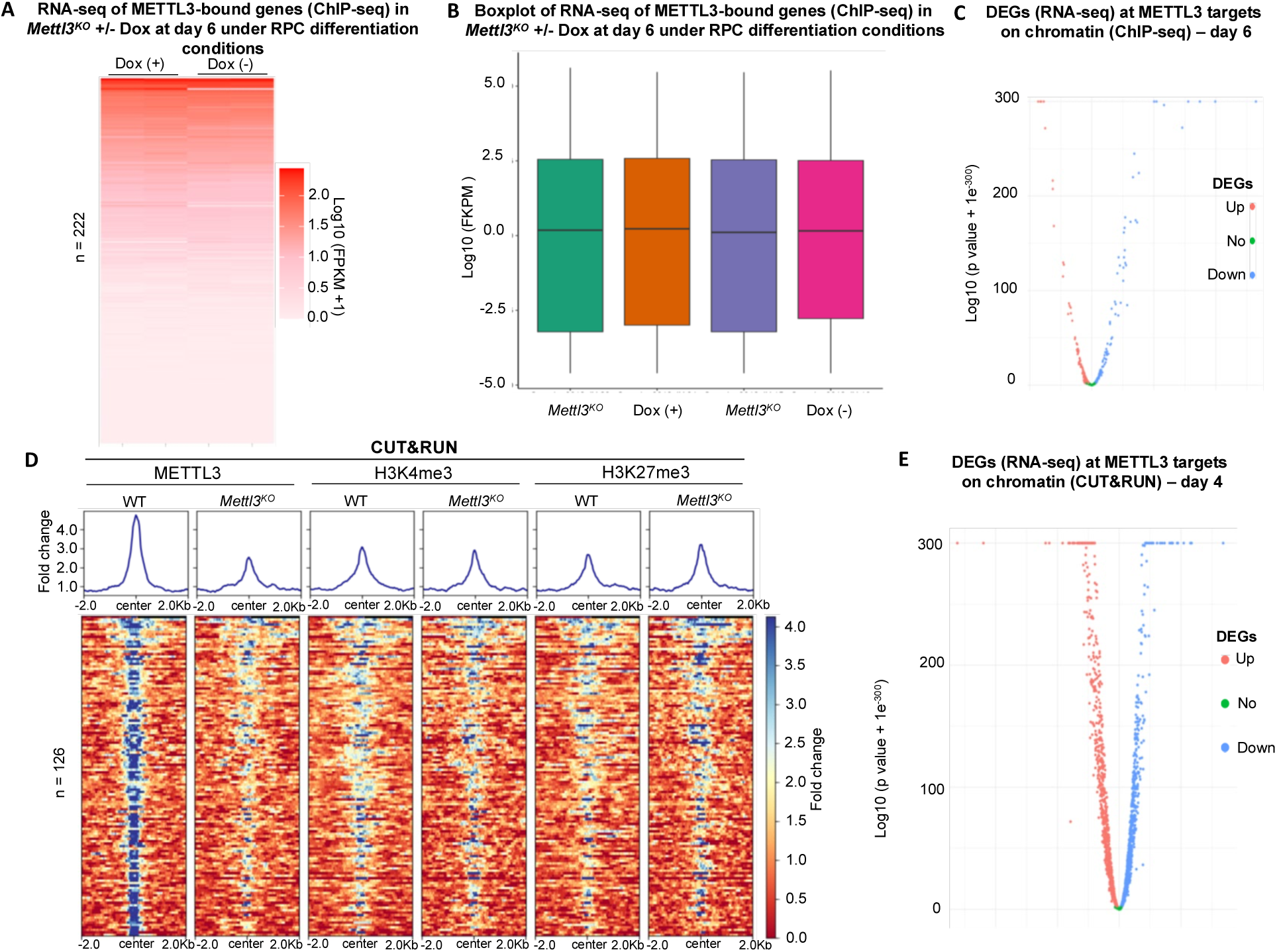
METTL3 regulates histone modifications but its direct targets on chromatin do not contribute to gene regulation during mESC-to-RPC differentiation. (**A, B**) RNA-seq heatmap and boxplot of METTL3-bound gene after 6 days under RPC differentiation culture of *Mettl3*^KO^ mESCs with or without Dox-rescue, n=3 independent experiments. (**C**) Volcano plot showed the differentially expressed genes (DEGs) from RNA-seq integrated with ChIP-seq from **A** and **B**. (**D**) H3K4me3 and H3K27me3 enrichment at METTL3-bound sites on chromatin in WT and *Mettl3*^KO^ organoids after 4 days under RPC differentiation conditions, n=3 independent experiments (**E**) Volcano plot showed the DEGs from RNA-seq integrated with CUT&RUN from **D**.

**Figure S3, related to Figure 3 and Discussion:**
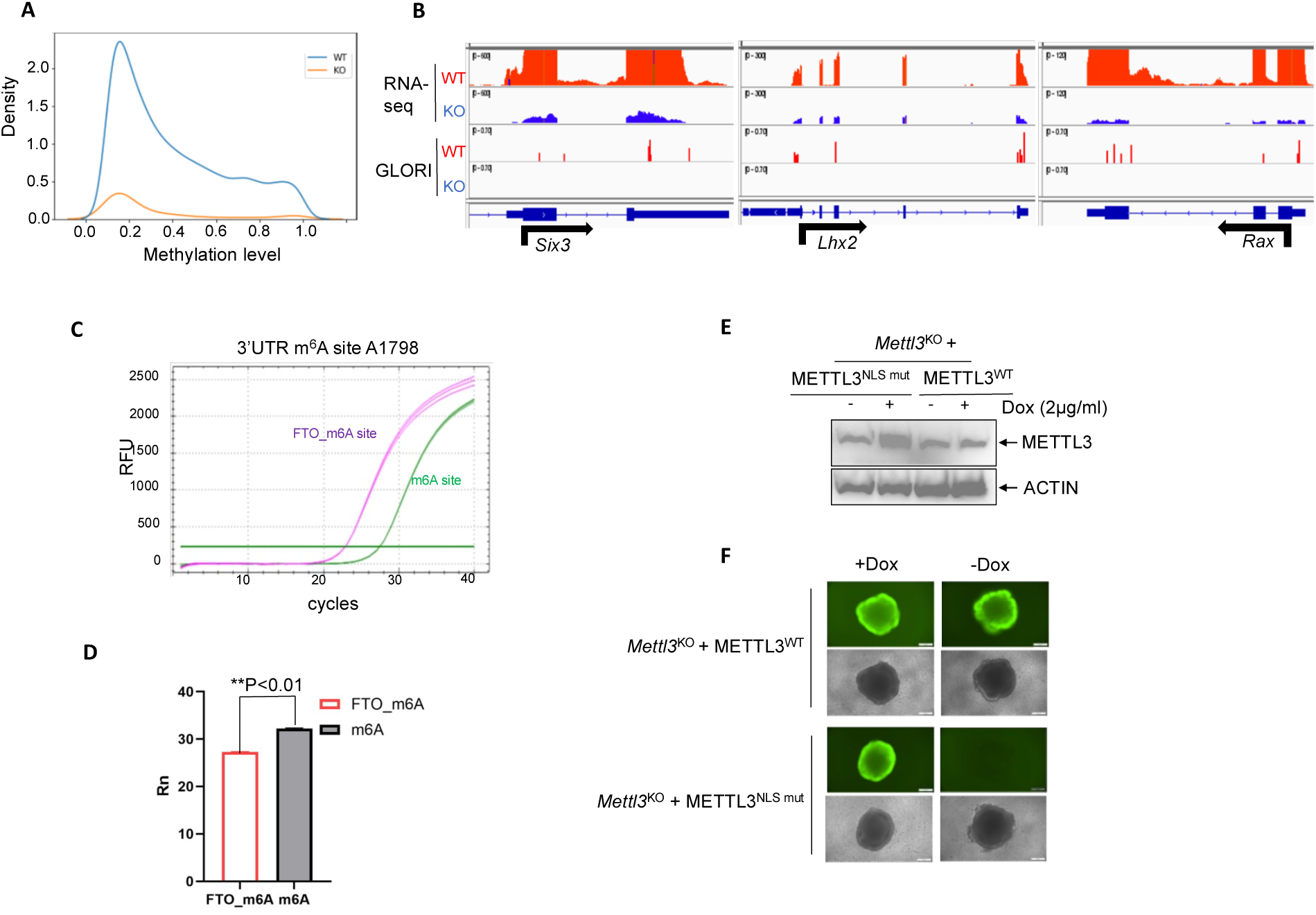
Function of METTL3-driven m6A modifications and transcriptional effects of acute METTL3 degradation in RPCs gene regulation during ESC to RPC differentiation. (**A**) Curve graph showed the m6A level distribution profiles in day 4 WT and *Mettl3^KO^* organoids centered on WT mRNAs with m6A marks, following 4 days under RPC differentiation conditions (vs Fig 3B, which shows total m6A sites detected by GLORI in each of WT and *Mettl3^KO^* organoid groups under identical RPC differentiation conditions). The median m6A modification level is ∼40%, n=3 independent experiments. (**B**) Track views of GLORI m6A sites (bottom two panels) aligned with RNA-seq (top two panels) for the *Mettl3^KO^* (KO) differentially regulated representative EFTF genes, *Six3*, *Lhx2 and Rax*. (**C, D**) SELECT assay showed that m6A removal at A1798 site of *Six3* 3’UTR dramatically increased *Six3* mRNA level. (**E,F**) The NLS mutant (mut) METTL3 could not rescue defective mESC-to-Rx:GFP^+^ RPC differentiation of *Mettl3^KO^* organoids. *Mettl3^KO^* mESCs were constitutively reconstituted with METTL3^NLS mut^ or METTL3^WT^ and differentiated into RPCs for 6 days treated with or without Dox (2μg/ml), which allows inducible expression of METTL3^WT^, n=3 independent experiments. (**E**) showed the protein levels of ectopically overexpressed METTL3^NLS mut^ or METTL3^WT^ in *Mettl3^KO^* Rx:GFP PRCs (day 6) by WB with METTL3 antibody detection (ACTIN: loading control). (**F**) Representative fluorescence microscopy images of *Mettl3^KO^* organoids reconstituted with METTL3^NLS mut^ or METTL3^WT^ and treated with or without Dox (2μg/ml). Scale bar: 200μm, n=3 independent experiments.

**Figure S4, related to Figure 4:**
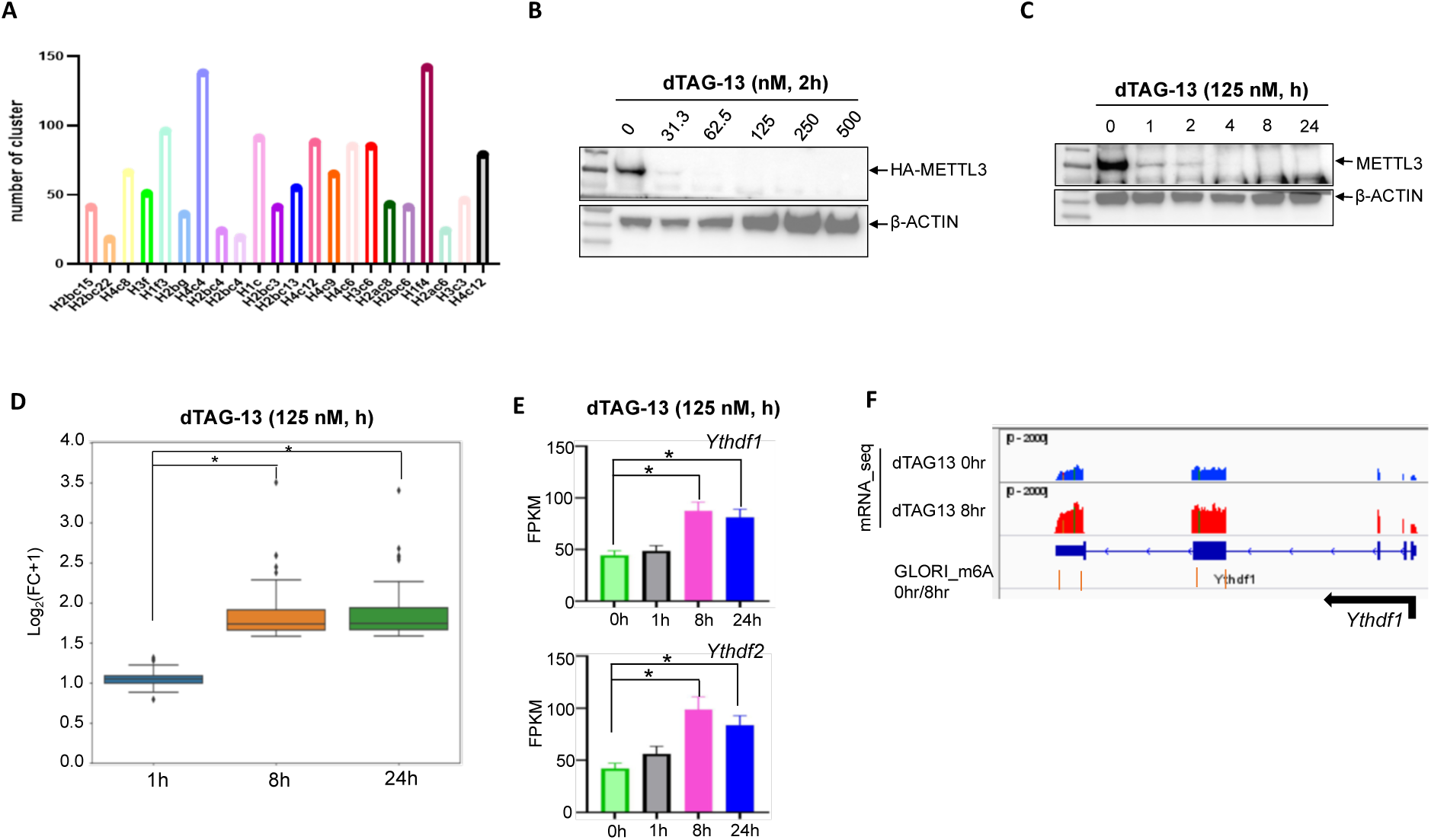
Acute METTL3 degradation reveals a regulatory METTL3-*Ythdf1* protein-RNA interaction and leads to impaired retinal differentiation. (**A**) METTL3 binds to histone RNA clusters as a quality-control for METTL3-eCLIP assay. (**B**, **C**) Acute degradation of endogenous METTL3 induced by dTAG-13 treatment. FKBP12^F36V^-HA was knocked into C-terminal region of METTL3 in Rx:GFP mESCs. At day 6 under RPC differentiation conditions, dose (B) and time (C) dependent degradation of FKBP12^F36V^-HA-METTL3 by dTAG-13 treatment were determined by Western blotting. HA and METTL3 antibodies were used separately for B and C, β-ACTIN as loading control. (**D**) Boxplot for up-regulated genes in day 6 RPCs at 8hr and 24hr post –METTL3 acute degradation, n=3 independent experiments. (**E**) *Ythdf1* and *Ythdf2* mRNA expression levels in day 6 RPCs upon acute METTL3 degradation at different time points, n=3 independent experiments. (**F**) Track views of GLORI m6A sites of *Ythdf1* mRNA upon acute METTL3 degradation by dTAG-13 (125nM, 0hr/8hr) at day 6 RPCs aligned with RNA-seq (top two panels).

**Supplemental Table 1:**
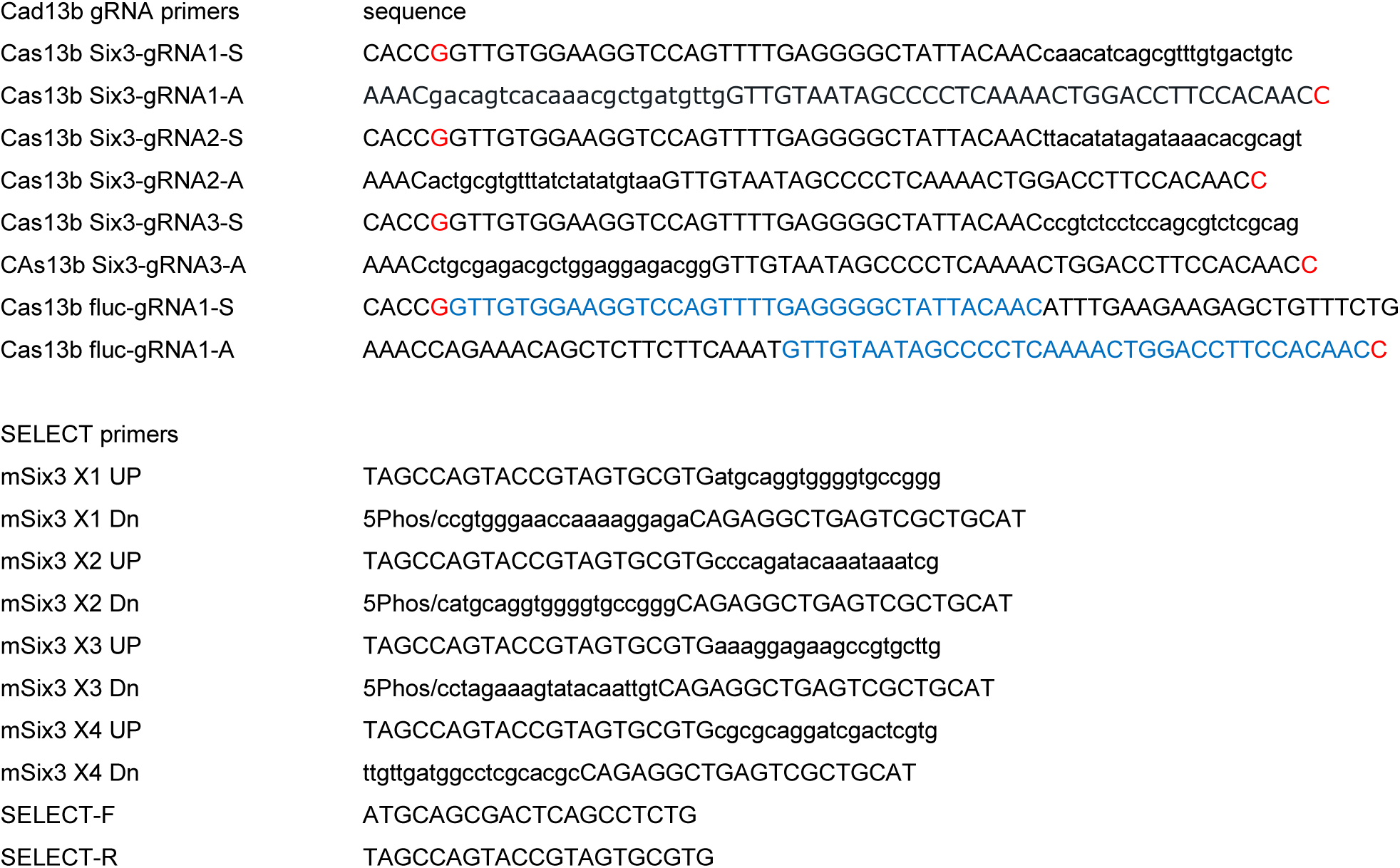
Cad13b gRNA primers and SELECT Primers.

## Notes

### Competing Interest Statement

The authors have declared no competing interest.

## References

1. Barbieri I, T. K., Pandolfini L, Shi J. … Promoter-bound METTL3 maintains myeloid leukaemia by m6A-dependent translation control Nature 2017, 552(7683): 126–131.

2. Deng, S., J. Zhang, J. Su … RNA m(6)A regulates transcription via DNA demethylation and chromatin accessibility Nat Genet 2022, 54(9): 1427–1437.

3. Dominissini D, M.-M. S., Schwartz S, Salmon-Divon M. … Topology of the human and mouse m6A RNA methylomes revealed by m6A-seq Nature 2012, 485: 201–206.

4. Eiraku M, T. N., Ishibashi H, Kawada M. … Self-organizing optic-cup morphogenesis in three-dimensional culture Nature 2011, 472(7341): 51–56.

5. Huang, H., Weng, H., Zhou, K. … Histone H3 trimethylation at lysine 36 guides m6A RNA modification co-transcriptionally Nature 2019, 567 414–419.

6. Huang L, L. H., Wang S, Chen S. … m6A writer complex promotes timely differentiation and survival of retinal progenitor cells in zebrafish Biochem Biophys Res Commun 2021, 567: 171–176.

7. Huang, X., J. Zhang, Y. Cun … Spatial control of m6A deposition on enhancer and promoter RNAs through co-acetylation of METTL3 and H3K27 on chromatin Mol Cell 2025, 85(7): 1349–1365.e1310.

8. Jin, K. X., R. Zuo, K. Anastassiadis … N6-methyladenosine (m(6)A) depletion regulates pluripotency exit by activating signaling pathways in embryonic stem cells Proc Natl Acad Sci U S A 2021, 118(51).

9. Khlebnikov, D. A., A. I. Nikolskaya, A. A. Zharikova … Comprehensive analysis of RNA-chromatin, RNA-, and DNA-protein interactions NAR Genom Bioinform 2025, 7(1): lqaf010.

10. Kim, K. L., P. van Galen, V. Hovestadt … Systematic detection of m(6)A-modified transcripts at single-molecule and single-cell resolution Cell Rep Methods 2021,1(5).

11. Knuckles P, C. S., Musheev M, Niehrs C. … RNA fate determination through cotranscriptional adenosine methylation and microprocessor binding Nat Struct Mol Biol 2017, 24(7): 561–569.

12. Lasman, L., V. Krupalnik, S. Viukov … Context-dependent functional compensation between Ythdf m(6)A reader proteins Genes Dev 2020, 34(19-20): 1373–1391.

13. Li, L., Y. Sun, A. E. Davis … Mettl14-mediated m(6)A modification ensures the cell-cycle progression of late-born retinal progenitor cells. Cell Rep 2023, 42(6): 112596.

14. Li, Q., Y. Huang, J. Xu … p53 inactivation unmasks histone methylation-independent WDR5 functions that drive self-renewal and differentiation of pluripotent stem cells Stem Cell Reports 2021, 16(11): 2642–2658.

15. Li, Q., F. Mao, B. Zhou … p53 Integrates Temporal WDR5 Inputs during Neuroectoderm and Mesoderm Differentiation of Mouse Embryonic Stem Cells Cell Rep 2020, 30(2): 465–480 e466.

16. Li Y, X. L., Tan K. … N6-Methyladenosine co-transcriptionally directs the demethylation of histone H3K9me2. Nat Genet. 2020, 52(9): 870–877.

17. Lin S, C. J., Du P, … The m(6)A Methyltransferase METTL3 Promotes Translation in Human Cancer Cells Mol Cell 2016, 62(3): 335–345.

18. Linder B, G. A., Olarerin-George AO. … Single-nucleotide-resolution mapping of m6A and m6Am throughout the transcriptome. Nat. Methods 2015, 12: 767–772.

19. Liu C, S. H., Yi Y. … Absolute quantification of single-base m6A methylation in the mammalian transcriptome using GLORI Nat Biotechnol 2023, 41(3): 355–366.

20. Liu, C., H. Sun, Y. Yi … Absolute quantification of single-base m(6)A methylation in the mammalian transcriptome using GLORI Nat Biotechnol 2023, 41(3): 355–366.

21. Liu, H., O. Begik, M. C. Lucas … Accurate detection of m(6)A RNA modifications in native RNA sequences Nat Commun 2019, 10(1): 4079.

22. Liu, J., X. Dou, C. Chen … N (6)-methyladenosine of chromosome-associated regulatory RNA regulates chromatin state and transcription Science 2020, 367(6477): 580–586.

23. Liu, J., Gao, M., He, J. … The RNA m6A reader YTHDC1 silences retrotransposons and guards ES cell identity Nature 2021, 591: 322–326.

24. Liu, J., Y. Yue, D. Han … A METTL3-METTL14 complex mediates mammalian nuclear RNA N6-adenosine methylation Nat Chem Biol 2014, 10(2): 93–95.

25. Liu P, L. F., Lin J. … m6A-independent genome-wide METTL3 and METTL14 redistribution drives the senescence-associated secretory phenotype Nat Cell Biol. 2021, 23(4): 355–365.

26. Liu X, W. C., Liu W. … Distinct features of H3K4me3 and H3K27me3 chromatin domains in pre-implantation embryos. Nature 2016, 537(7621): 558–562.

27. Meyer KD, S. Y., Zumbo P. … Comprehensive analysis of mRNA methylation reveals enrichment in 3′ UTRs and near stop codons Cell 2012, 149: 1635–1646

28. Murakami, K., U. Gunesdogan, J. J. Zylicz … NANOG alone induces germ cells in primed epiblast in vitro by activation of enhancers Nature 2016, 529(7586): 403–407.

29. Nabet B, R. J., Buckley DL, Paulk J. … The dTAG system for immediate and target-specific protein degradation Nat Chem Biol. 2018, 14(5): 431–441.

30. Niu, F., P. Che, Z. Yang … m(6)A regulation of cortical and retinal neurogenesis is mediated by the redundant m(6)A readers YTHDFs iScience 2022, 25(9): 104908.

31. Quarto, G., A. Li Greci, M. Bizet … Fine-tuning of gene expression through the Mettl3-Mettl14-Dnmt1 axis controls ESC differentiation Cell 2025, 188(4): 998–1018.e1026.

32. Ruthenburg AJ, A. C., Wysocka J. … Methylation of lysine 4 on histone H3: intricacy of writing and reading a single epigenetic mark. Mol Cell 2007, 25(1): 15–30.

33. Schöller E, W. F., Treiber T. … Interactions, localization, and phosphorylation of the m6A generating METTL3-METTL14-WTAP complex RNA 2018, 24(4): 499–512.

34. Schwartz YB, P. V. Polycomb silencing mechanisms and the management of genomic programmes Nat Rev Genet. 2007, 8(1): 9–22.

35. Shi H, W. X., Lu Z. … YTHDF3 facilitates translation and decay of N6-methyladenosine-modified RNA Cell Res. 2017, 27(3): 315–328.

36. Van Nostrand EL, P. G., Shishkin AA, Gelboin-Burkhart C. … Robust transcriptome-wide discovery of RNA-binding protein binding sites with enhanced CLIP (eCLIP) Nat Methods 2016, 13(6): 508–514.

37. Wang P, D. K., Nam Y. … Structural Basis for Cooperative Function of Mettl3 and Mettl14 Methyltransferases Mol Cell 2016, 63(2): 306–317.

38. Wang, X., Z. Lu, A. Gomez … N6-methyladenosine-dependent regulation of messenger RNA stability Nature 2014, 505(7481): 117–120.

39. Wang X, Z. B., Roundtree IA. … N(6)-methyladenosine Modulates Messenger RNA Translation Efficiency Cell 2015, 161(6): 1388–1399.

40. Wilinski, D., Dus, M. … N6-adenosine methylation controls the translation of insulin mRNA Nat Struct Mol Biol 2023, 30: 1260–1264.

41. Xiao, Y., Y. Wang, Q. Tang … An Elongation- and Ligation-Based qPCR Amplification Method for the Radiolabeling-Free Detection of Locus-Specific N(6)-Methyladenosine Modification Angew Chem Int Ed Engl 2018, 57(49): 15995–16000.

42. Xin, Y., Q. He, H. … m(6)A epitranscriptomic modification regulates neural progenitor-to-glial cell transition in the retina Elife 2022, 11:e79994.

43. Xu, W., J. Li, C. He … METTL3 regulates heterochromatin in mouse embryonic stem cells Nature 2021, 591(7849): 317–321.

44. Zaccara S, J. S. A Unified Model for the Function of YTHDF Proteins in Regulating m6A-Modified mRNA Cell 2020, 181(7): 1582–1595.

45. Zaccara S, J. S. (Understanding the redundant functions of the m6A-binding YTHDF proteins RNA 2024, 30(5): 468–481.

